# Isolation of rhizobia from Ontario soils that are effective at fixing nitrogen with common bean (*Phaseolus vulgaris*)

**DOI:** 10.64898/2026.05.01.722220

**Authors:** Tia L. Harrison, Upkardeep S. Pandher, Avery Dixon, Oona Esme, Erika M.H. Gagnon, Natalia Naranjo-Robayo, Rebecca T. Doyle, Ivan J. Oresnik, George C. diCenzo

## Abstract

Common bean (*Phaseolus vulgaris*) is an important crop in Canada and globally. Like other legumes, common bean (*Phaseolus vulgaris*) establishes symbiotic interactions with nitrogen fixing bacteria called rhizobia. However, nitrogen fixation by rhizobia in association with common bean is often suboptimal, constraining its productivity and necessitating the application of nitrogen fertilizer. To support the development of high-performing, locally adapted rhizobial inoculants for Ontario common bean growers, we isolated 216 common bean-nodulating rhizobia from southern Ontario soils using a nodule trapping approach with four common bean cultivars. Whole genome sequencing followed by phylogenomic analyses of the 216 rhizobial isolates revealed substantial diversity, assigning them to 11 *Rhizobium* species, including two novel species. Nearly all isolates belong to the symbiovar *phaseoli*, spanning the *nodC* γ-a, γ-b, and α alleles, with four isolates belonging to the symbiovar *gallica*. Soil origin had a significant impact on the species-level community composition recovered during the nodule trapping experiments, indicative of biogeographical structuring of common bean-nodulating rhizobia across southern Ontario. In contrast, host trapping cultivar had only a minor influence of the recovered *Rhizobium* population diversity. Greenhouse assays demonstrated that one of the novel *Rhizobium* species exhibited the highest average symbiotic effectiveness, although high-quality isolates were found across multiple species. Together, these results revealed a diverse and genomically variable *Rhizobium* community capable of forming effective symbioses with common bean in southern Ontario soils. Importantly, our genome-sequenced *Rhizobium* collection will serve as a valuable resource for identifying competitive and high-quality strains for the development of inoculants tailored to Ontario common bean production.

**IMPORTANCE:** Common bean is a globally important food crop, yet its productivity is often limited by suboptimal nitrogen fixation, forcing growers to rely on synthetic fertilizers. Consequently, identifying high-performing, locally adapted inoculant strains is essential for reducing dependence on synthetic nitrogen fertilizers and improving the sustainability of temperate agroecosystems. Our study provides a genome-sequenced collection of common bean–nodulating *Rhizobium* from southern Ontario, revealing substantial species and genomic diversity across sampling locations. Greenhouse studies allowed us to identify multiple isolates, including isolates from a novel *Rhizobium* species, that consistently fix nitrogen with, and enhance the growth of, common bean plants. Our findings highlight strong biogeographical structuring of rhizobial communities and demonstrate that Ontario soils already harbour strains with high symbiotic potential. In addition, our *Rhizobium* collection represents a foundational resource to support future inoculant development and enables future work on the ecology, evolution, and applied optimization of legume–rhizobium symbioses.

## INTRODUCTION

Common bean (*Phaseolus vulgaris*) is a globally cultivated crop that plays a crucial role in human nutrition and local economies as they are an affordable source of protein, fibre, and minerals (1). Wild common bean likely originated in Mexico and then spread throughout Central and South America, leading to the emergence of three distinct gene pools in Mesoamerica, the Andes, and northern Peru–Ecuador (2–4). Domestication of common bean is estimated to have occurred independently in both Mesoamerica and the Andes ∼ 8,000 years ago (5). Cultivation of diverse varieties of common bean has since spread globally, being grown both as a vegetable (e.g., green and yellow string beans) and as a pulse crop (e.g., navy beans, kidney bean, black bean, and pinto bean). When grown as a pulse, common bean is often referred to as “dry edible bean” or just “dry bean”.

Like other legumes, common bean can form a nitrogen-fixing symbiosis with rhizobia. Rhizobia are a polyphyletic group of soil bacteria of the classes *Alphaproteobacteria* and *Betaproteobacteria* that, following an exchange of signals with a compatible legume host, intracellularly colonize a specialized legume organ known as a root nodule (6–8). Within the nodule, the rhizobia convert atmospheric nitrogen into ammonia, which is provided to their plant host in exchange for carbon (9). Although common bean can form nitrogen-fixing symbioses with rhizobia, it is often considered a “poor nitrogen-fixer” (note that the nitrogen-fixation is done by the rhizobia, not the plant) compared to other legume species including soybean (*Glycine max*) and faba bean (*Vicia faba*) (10, 11). The low rates of nitrogen fixation in common bean nodules may be partially due to its broad symbiont breadth, forming nodules with a broad range of rhizobial species, including those from the bacterial genera *Rhizobium*, *Pararhizobium*, *Sinorhizobium*, *Bradyrhizobium*, *Paraburkholderia*, and *Cupriavidus* (11, 12). However, many of the rhizobia that interact with common bean may be poorly adapted to fix nitrogen with this plant species (12), resulting in the frequent formation of suboptimal associations between common bean and its rhizobial partners.

The poor nitrogen-fixing symbiosis characteristic of common bean has significant agricultural impacts. For many legume crops, growers have access to commercially available rhizobial inoculants, which are products that are applied to seeds or fields and contain elite rhizobium strains compatible with the legume species. High-quality rhizobial inoculants, those that effectively fix nitrogen, are available for numerous legume crops globally, and in many cases (e.g., soybean), the amount of fixed nitrogen these inoculants provide is sufficient to replace the need for nitrogen fertilizer, making their use economically viable (13, 14). On the other hand, inoculants developed for common bean worldwide have shown variable levels of success (15–17). Consequently, in many jurisdictions, growers are unable to rely solely on nitrogen fixation to support common bean yields, and instead, must apply chemical nitrogen fertilizers. Although nitrogen fertilizers are important to support crop yields and food security (18), they are expensive for farmers to purchase and have negative impacts on the environment, including the use of fossil fuels in their production and conversion of a portion of the nitrogen to the potent greenhouse gas nitrous oxide (19, 20).

Common bean is an important crop in Canada, particularly in Ontario and Manitoba which together accounted for over 75% of national production between 2023 and 2024 (21). Canada is the 16^th^ largest producer of dry bean globally, and one of the top five exporters, producing and exporting around 390,000 and 360,000 tonnes (valued at ∼370 million USD), respectively, per year from 2020-2022 (21). Common bean crops in Canada are generally not inoculated, and instead, are fertilized with ∼40 kg/ha of nitrogen (22), although some data suggest this application may not be required to achieve high yields (23). Improving the capacity for nitrogen-fixing symbiosis with Canadian common bean crops therefore has the potential to reduce input costs for Canadian growers and reduce the climate impact of common bean cultivation. Tackling this challenge will likely involve breeding common bean for varieties with higher capacity to support nitrogen-fixing symbiosis (24, 25), in conjunction with developing high-quality rhizobial inoculants specific for this crop species. Studies have repeatedly shown that inoculants developed from locally sourced rhizobia outperform inoculants developed from rhizobia sourced elsewhere (26–30), likely due to rhizobial adaptation to local soil and climatic conditions as well as compatibility with the existing soil microbiome (31). Consequently, developing commercial rhizobial inoculants for common bean growers in Canada requires building a collection of locally sourced isolates adapted to Canadian agro-climatic conditions.

Here, we report a genome-sequenced collection of 216 rhizobial isolates capable of nodulating common bean. These rhizobia were isolated from seven sites collected across southern Ontario using a nodule trapping approach, which resulted in a genetically diverse collection spanning 11 species of the genus *Rhizobium*. We found that the location of the source soils influenced which *Rhizobium* species were recovered, whereas there were only minor impacts of common bean cultivar. Phenotypic characterization of the isolates revealed extensive variation in nitrogen fixation abilities when in symbiosis with common bean grown in greenhouse experiments. Overall, our data demonstrate that

Ontario soils contain diverse rhizobia capable of fixing nitrogen with common bean, with one of two novel *Rhizobium* species identified being a higher-quality symbiont on average compared to the other *Rhizobium* species.

## METHODS

### Statistical analyses and data visualization

Statistical analyses and data visualization were performed in RStudio version 2025.9.2.418 (32) with R version 4.5.2 (33) and the packages canadianmaps version 2.0.0 (34), car version 3.1-3 (35), dplyr version 1.1.4 (36), emmeans version 2.0.1 (37), gggenes version 0.6.0 (38), ggplot2 version 4.0.1 (39), gplots version 3.3.0 (40), knitr version 1.51 (41), lme4 version 1.1-38 (42), multcomp version 1.4-29 (43), patchwork version 1.3.2 (44), phyloseq version 1.54 (45), rcartocolor version 2.1.2 (46), RColourBrewer version 1.1-3 (47), sf version 1.1-0 (48), tibble version 3.3.1 (49), tidyr version 1.3.2 (50), tidyverse version 2.0.0 (51), UpSetR version 1.4.0 (52), and vegan version 2.7-2 (53).

### Soil collection

Thirteen soil samples from six sites were collected from southwestern to southeastern Ontario (**Table 1, Figure S1**) from May to June, 2023. Four samples were collected from the Huron Research Station (Exeter, Ontario) that has a long history of bean cultivation; two of these samples were collected within 10 m of each other from a field sowed with *P. vulgaris* the previous year, while the other two were collected within 10 m of each other from a field sowed with *Vigna angularis* (adzuki bean) the previous year. The four samples from the Huron Research Station were kept separate for downstream processing. An additional sample came from a commercial farm in Elgin, Ontario, which also had a long history of *P. vulgaris* cultivation, although it was last seeded with *P. vulgaris* in 2020. Three samples were collected from a community garden in Kingston, Ontario, each from separate plots that had been sowed with a different heritage variety of *P. vulgaris* the previous year; these samples were combined prior to downstream processing. One sample was collected from a home garden in Kingston, Ontario, which was seeded with *P. vulgaris* the previous season. One sample was collected from a site of a previous home garden in Brampton, Ontario, which had no known history of *P. vulgaris* cultivation. Lastly, three samples were collected from a home garden in Caledon, Ontario. Two of these plots were sowed with *P. vulgaris* the previous year while the third was sowed with *Vicia faba* (faba bean); these samples were combined prior to downstream processing. Following the combination of related soil samples, nine soil samples remained for subsequent rhizobial isolation.

**Table 1.**
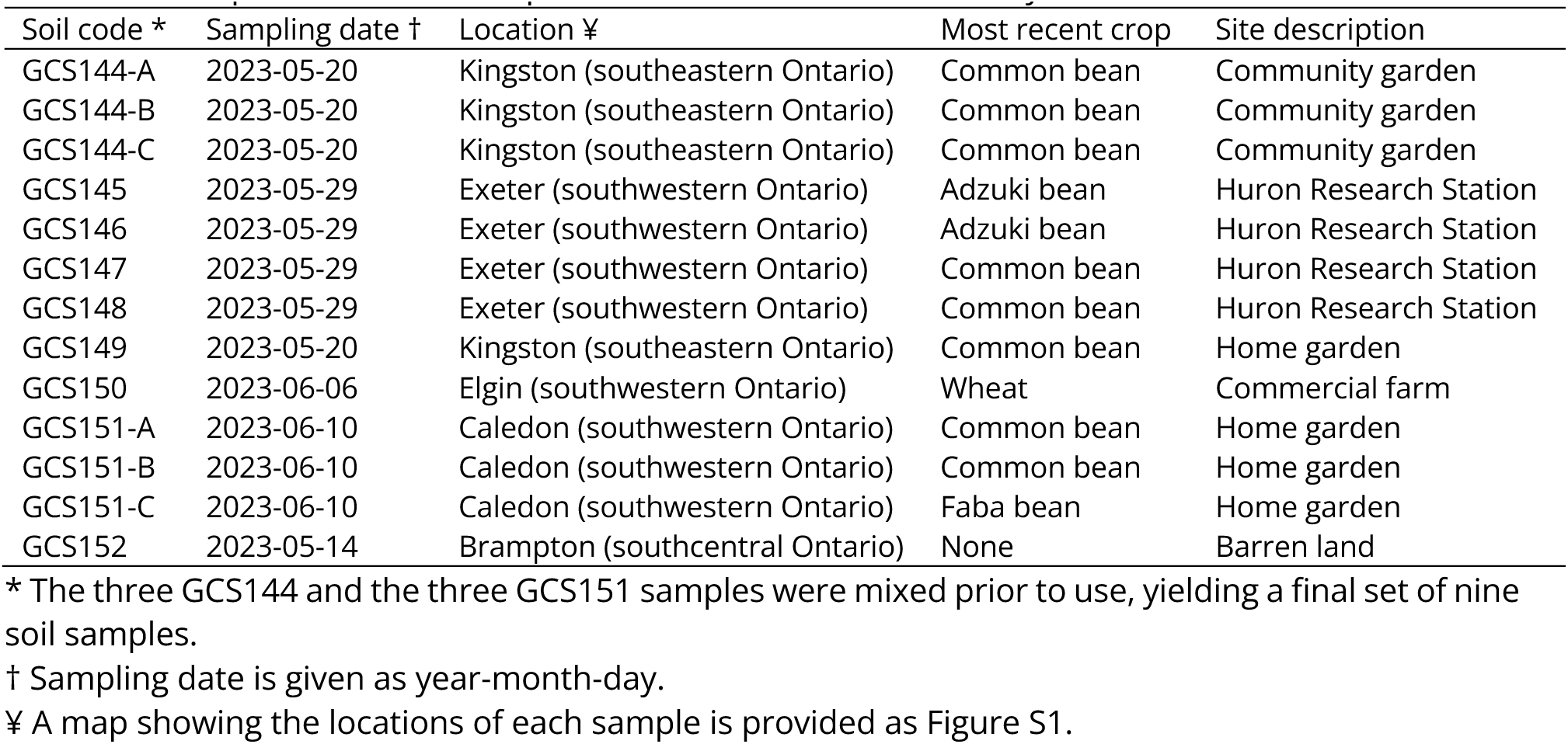
Description of 13 soil samples and the sites from which they were collected.

| Soil code * | Sampling date † | Location ¥ | Most recent crop | Site description |
| --- | --- | --- | --- | --- |
| GCS144-A | 2023-05-20 | Kingston (southeastern Ontario) | Common bean | Community garden |
| GCS144-B | 2023-05-20 | Kingston (southeastern Ontario) | Common bean | Community garden |
| GCS144-C | 2023-05-20 | Kingston (southeastern Ontario) | Common bean | Community garden |
| GCS145 | 2023-05-29 | Exeter (southwestern Ontario) | Adzuki bean | Huron Research Station |
| GCS146 | 2023-05-29 | Exeter (southwestern Ontario) | Adzuki bean | Huron Research Station |
| GCS147 | 2023-05-29 | Exeter (southwestern Ontario) | Common bean | Huron Research Station |
| GCS148 | 2023-05-29 | Exeter (southwestern Ontario) | Common bean | Huron Research Station |
| GCS149 | 2023-05-20 | Kingston (southeastern Ontario) | Common bean | Home garden |
| GCS150 | 2023-06-06 | Elgin (southwestern Ontario) | Wheat | Commercial farm |
| GCS151-A | 2023-06-10 | Caledon (southwestern Ontario) | Common bean | Home garden |
| GCS151-B | 2023-06-10 | Caledon (southwestern Ontario) | Common bean | Home garden |
| GCS151-C | 2023-06-10 | Caledon (southwestern Ontario) | Faba bean | Home garden |
| GCS152 | 2023-05-14 | Brampton (southcentral Ontario) | None | Barren land |

\* The three GCS144 and the three GCS151 samples were mixed prior to use, yielding a final set of nine soil samples.
† Sampling date is given as year-month-day.
¥ A map showing the locations of each sample is provided as Figure S1.

### Plant material

Four varieties of *P. vulgaris* were used for nodule trapping experiments: a commercial variety of navy bean (AAC Shock), a commercial variety of dark red kidney bean (Dynasty), an unnamed heritage navy bean variety, and an unnamed heritage kidney bean variety. AAC Shock was used for all subsequent experiments that phenotypically characterized rhizobial isolates.

### Bacterial growth conditions

Rhizobia were grown at 28°C using either tryptone – yeast extract (TY) medium (5 g/L tryptone, 2.5 g/L yeast extract, 10 mmol/L calcium cloride, and 15 g/L agar for solid medium) or yeast extract – mannitol (YM) medium (5 g/L mannitol, 0.5 g/L dipotassium phosphate, 0.1 g/L sodium chloride, 0.2 g/L magnesium sulfate heptahydrate, 0.4 g/L yeast extract, and 15 g/L agar for solid medium).

### Plant growth conditions

*P. vulgaris* seeds were surface sterilized with 95% ethanol for 30 seconds, rinsed twice with sterile ddH_2_O, soaked in ∼1% sodium hypochlorite for 3 min, and rinsed two to five times (until the bleach odour was no longer detectable) with sterile ddH_2_O for 2 min per rinse. Sterilized seeds were spread on sterile water agar (1% agar in water) plates and left to germinate at room temperature for three days in the dark. Seedlings were planted in autoclaved Leonard Assemblies, which consisted of two stacked Magenta Jars connected by a cotton wick that extended from the top jar (containing a 1:1 (w/w) mixture of vermiculite and silica sand) to the bottom jar (containing 250 mL of Jensen’s medium that was initially poured over the top jar) (54, 55). One or two seedlings were planted per Leonard Assembly, and when necessary, Jensen’s medium was supplemented with nitrogen as described below. The plants were grown in the Queen’s University Phytotron’s greenhouse for two days, at which point they were inoculated with soil slurries (nodule trapping) or rhizobial cultures (phenotypic characterization), as described below. Plants were then grown for four to five weeks in the Queen’s University Phytotron’s greenhouse. Plants were routinely watered with sterile ddH_2_O and watered once with 250 mL of nitrogen-free Jensen’s medium 21 days post-inoculation.

### Nodule trapping experiments

Nodule trapping experiments were used to isolate rhizobia from each of the nine final soil samples. Soil slurries were prepared by adding a sufficient volume of sterile ddH_2_O to 5 g of soil to produce a 50 mL soil slurry, after which 5 mL aliquots were mixed with 20 mL of sterile ddH_2_O. We then inoculated each of the four common bean varieties in Leonard Assemblies (one seedling per assembly) with each of the nine soils yielding a total of 36 plants. A 37^th^ pot was accidentally inoculated with two slurries; we still sampled nodules from this pot and include the isolates in our collection, but we excluded these isolates from our nodule trapping statistical analysis. After four weeks, we harvested all plants and collected the six largest and most pink (an indication of high nitrogen fixation) nodules per plant. Nodules were rinsed with 0.85% NaCl, surface sterilized with 1% sodium hypochlorite for 10 minutes, and then rinsed with YM broth. Nodules were then crushed in 500 µl of YM broth with a sterile pestle in 1.5 mL centrifuge tubes. A total of 10 µl of the resulting solution was plated on YM agar plates and incubated at 28°C for three to five days. Rhizobia were streak purified twice on YM agar and then once on TY agar to generate pure lineages. In some cases, if more than one colony morphology was observed, one colony from each morphology type was streak purified from the same agar plate. Finally, purified isolates were archived in TY broth containing 7% dimethyl sulfoxide at -80°C as part of the Canadian Collection of Agricultural Soil Microbes. In total, 216 rhizobia and 11 non-rhizobia were archived (**Dataset S1**); we only focus on the rhizobia in this manuscript.

To examine the impact of source soil and host genotype on the species-level composition of the rhizobial populations recovered during the nodule trapping experiments, an R phyloseq object was created that included species counts, the taxonomic classification of each species, source soil, and host genotype; each soil – host genotype pairing was treated as a separate sample. Species count data was then converted to relative abundance by dividing each count by the total counts in that sample. Bray-Curtis distances were calculated using the distance function in phyloseq. The betadisper and permutest functions of the vegan package were used to test if the within-group dispersion per treatment was equal, while a PERMANOVA was run using the adonis2 function of vegan. A Principal Component Analysis (PCoA) and a capscale analysis were run using the ordinate function of phyloseq, and the results plotted using the plot_ordination function of phyloseq. The PERMANOVA and capscale analyses were run with the distance matrix as the response variable and with soil and host genotype as the main effects.

### Preliminary screening of all *Rhizobium* isolates for nitrogen-fixation capacity

In November 2023, a greenhouse experiment was conducted to screen all 216 *Rhizobium* isolates for their ability to fix nitrogen in symbiosis with common bean. *Rhizobium* isolates were grown in 1 mL of TY broth in 24-well microtiter plates for two days and then diluted by mixing 0.9 mL of culture with 4.1 mL of sterile ddH_2_O. One mL of the resulting diluted culture for each strain was then inoculated into Leonard Assemblies containing two seedlings each. In addition to inoculated plants, 15 Leonard Assemblies were left uninoculated, eight for nitrogen-plus controls (which were watered with 20 mM potassium nitrate every 15 days) and seven for nitrogen-minus controls (which did not receive potassium nitrate). In total, 231 plants (216 inoculated + 15 uninoculated controls) were left to grow in the Queen’s University Phytotron’s greenhouse for five weeks, with assemblies thinned to one live plant per pot four days post-inoculation. Starting seven days post-inoculation and continuing weekly until the end of the experiment, a PlantPen NDVI 300 handheld device was used to measure the Normalized Difference Vegetation Index (NDVI) of the newest leaf on each inoculated plant, which reflects leaf chlorophyll A content indicative of nitrogen fixation. After five weeks of growth, all plants were harvested, and plant shoots were collected and dried at 60°C for three weeks and then weighed. Linear models were fit with shoot dry weight as the response variable, rhizobium species (11 species total) as the explanatory predictor, and added offsets to the model to account for the unequal sample sizes of the groups. The raw data met model assumptions of linearity, normality, and homoscedasticity, and so no data transformations were applied. Significance was tested using a Type II ANOVA. Pairwise differences between groups were assessed using estimated marginal means (emmeans) with Tukey’s adjustment for multiple comparisons. A linear regression was also conducted to test whether NDVI values recorded at the end of week 3 were predictive of final shoot dry weights measured at the end of week 5, with significance tested using a Type II ANOVA.

### Evaluating the impact of nitrogen supplementation on *Rhizobium* inoculation

From April to May, 2024, a second greenhouse experiment was conducted to examine potential synergistic effects of *Rhizobium* inoculation and nitrogen addition on the growth of common bean. Two types of nitrogen regimens were tested: one-time application of “starter” nitrogen (potassium nitrate) at concentrations of 0 mM, 2.5 mM, 5 mM, 10 mM, and 20 mM, or ongoing application of nitrogen (potassium nitrate) at concentrations of 1 mM, 2.5 mM, 10 mM, and 20 mM. Leonard Assemblies (one seedling per assembly) were prepared as described above, except that the Jensen’s medium was supplemented with potassium nitrate at the required concentrations. *Rhizobium* sp. QUR0071 was grown in TY broth for two days and then diluted 1:5 with sterile ddH_2_O, resulting in a final optical density at 600 nm (OD600) of 0.208. One mL of the diluted culture was then used to inoculate seedlings in half of the Leonard Assemblies. Triplicate Leonard Assemblies were prepared for each condition, yielding 54 assemblies (nine nitrogen regimens by two inoculation conditions [with or without rhizobium], each in triplicate). Plants were grown in the Queen’s University Phytotron’s greenhouse for four weeks. Plants from the ongoing application of nitrogen treatments were watered with potassium nitrate at the appropriate concentrations every eight days, whereas all other plants received no nitrogen during growth. NDVI was measured for the top newest leaf and bottom-most leaf at weeks 3 and 4. After four weeks of growth, all plants were harvested, and plant shoots were collected and dried at 60°C for one week and then weighed.

Data collected from the ongoing nitrogen treatments and one time nitrogen treatments were analyzed separately. In both cases, linear models were fit with shoot mass as the response variable, and included rhizobium presence/absence, nitrogen concentration (treated as an ordered factor rather than a continuous variable), and their interaction as explanatory predictors. shoot biomass data was log transformed to meet assumptions of linearity. Significance was tested using Type III ANOVAs, while pairwise contrasts between treatments with and without rhizobia at a given nitrogen concentration were assessed using emmeans with Tukey’s adjustment for multiple comparisons.

### Repeat screening of a subset of *Rhizobium* isolates for nitrogen-fixation capacity

In September 2025, a greenhouse experiment was conducted at Queen’s University to test the impacts of a subset of 12 rhizobial isolates on common bean growth, using a randomized block experimental design. Leonard Assemblies were prepared containing two seedlings per assembly as described above except that the Jensen’s medium was supplemented with 5 mM potassium nitrate based on the results of the previous experiment. Rhizobial isolates were grown in TY broth for two days and then diluted to an OD600 of 0.1 in 0.85% NaCl. One mL of each diluted culture was individually mixed with 10 mL of sterile ddH_2_O and used to inoculate the seedlings in one Leonard Assembly; five replicate Leonard Assemblies were prepared per isolate, and five uninoculated controls were also prepared. Plants were left to grow in the Queen’s University Phytotron’s greenhouse for four weeks and thinned to one seedling per pot three days post-inoculation. NDVI was measured on the top newest leaf of each plant weekly starting two weeks post-inoculation and plants were harvested for shoot dry weight measurements four weeks post-inoculation.

Data were analyzed in R using a linear mixed-effect model that included shoot dry weight as the response variable, strain as the fixed effect (12 groups), and block as a random factor (five groups). All model assumptions of linearity, normality, and homoscedasticity were met with raw shoot mass data values. Significance was tested using a Type II ANOVA, while pairwise contrasts between strains were assessed using emmeans with Tukey’s adjustment for multiple comparisons.

### Whole genome sequencing

Single colonies were inoculated into TY broth and grown for up to four days, following which DNA was extracted using DNeasy UltraClean 96 Microbial Kits (Qiagen) according to the manufacturer’s instructions with the following modifications. All centrifugation steps were performed at a speed of 3,500 x *g*, 125 µL of a 20 ng/µL RNase A solution (New England Biolabs; Catalog No. T3018L) solution was added to 15 mL of SL buffer prior to lysis, and all samples were eluted using sterile ddH_2_O. Finally, the DNA was eluted from the columns using 100 µL of nuclease-free water. Subsequently, Oxford Nanopore Technologies (ONT) library preparation was performed using a Rapid Barcoding Kit 96 V14 (SQK-RBK114.96, ONT) according to the manufacturer’s instructions, followed by sequencing on PromethION flow cells (R10.4.1, ONT) on a P2 Solo device. Basecalling and demultiplexing were performed using dorado version 0.4.1 with the model dna_r10.4.1_e8.2_400bps_sup@v4.2.0.

### Genome assembly, annotation, and taxonomic classification

Genome assembly was performed using the raw ONT reads with Flye version 2.9.3 (56), followed by polishing using the ONT reads with Medaka version 1.10.0 (github.com/nanoporetech/medaka). Next, Pullseq version 1.0.2 (github.com/bcthomas/pullseq) was used to remove contigs less than 5,000 bp in length. Genome assembly quality was determined using CheckM version 1.2.2 (57) and initial taxonomic classification performed with the Genome Taxonomy Database Toolkit (GTDB-Tk) version 2.3.0 with database version R214 (58). Genome annotation was then performed using the NCBI Prokaryotic Genome Annotation Pipeline (PGAP) version input-2023-10-03.build7061 (59). Lastly, the taxonomic classification of each *Rhizobium* isolate was assigned by comparing each isolate’s genome to a *Rhizobium* type strain reference library (60) using FastANI version 1.33 (61). When the average nucleotide identity (ANI) compared to the top hit was ≥ 95%, the isolate was assigned to the same species as the top hit. Otherwise, the isolate was not assigned a species level classification; these isolates were then grouped into putatively novel species using an ANI threshold of 95%.

### Species phylogenomic analyses

ANI calculations between each pair of *Rhizobium* isolates from our study were performed using FastANI. Subsequently, a representative subset of 37 isolates was selected by dereplicating the dataset using dRep version 3.5.0 (62) with a secondary clustering threshold of 99.8%. We additionally downloaded the published genomes for all 47 *Rhizobium* type strains and three type strains from the genus *Martinezella*, which were downloaded from the National Center for Biotechnology Information (NCBI) database (**Dataset S2**). These 87 genomes (37 representative isolates from our collection plus the 50 type strains) were then used to construct a core-proteome phylogeny using an established pipeline (github.com/diCenzo-Lab/017_2025_Rhizobiaceae_taxonomy) as described previously (60, 63). Briefly, TBLASTN version 2.17.0 (64) was used to identify orthologs of 170 conserved, single-copy, non-recombining genes in each genome. The corresponding protein sequences were then aligned with MAFFT version 7.453 (65) and trimmed using trimAl version 1.4.rev22 with the automated1 algorithm (66), after which the alignments were concatenated. The concatenated alignment was then used as input for IQ-TREE version 2.2.2.4 (67) with the substitution model LG+F+I+R5, as ModelFinder (68) indicated this was the best-scoring model. Branch support was assessed using the Shimodaira–Hasegawa-like approximate likelihood ratio test (SH-aLRT) and ultrafast jackknife analysis set to a subsampling proportion of 40%, with both metrics calculated from 1,000 replicates. Phylogenies were visualized using the iTOL webserver (69). Lastly, linear models with genome size or contig number as the response variable and species as the main effect, were run to test whether the species differed in genome size and/or number of replicons.

### Phylogenetic analyses of NodC

NodC homologs were extracted from the proteomes of the 37 dereplicated *Rhizobium* isolates using a previously described pipeline (70). Briefly, the proteomes of all 37 isolates were first searched using a hidden Markov model (HMM) built from the TIGRFAM NodC family (TIGR04245) using HMMER version 3.3 (71). All hits were extracted and then compared to a HMM database consisting of all HMMs from the Pfam version 37.0 (72) and TIGRFAM version 15.0 (73) databases; proteins whose top HMM hit was TIGR04245 were annotated as NodC. Subsequently, the nucleotide sequences of the corresponding genes were extracted and MAFFT version 7.471 (65) used to create a multi-sequence alignment of these 37 *nodC* sequences together with the 54 *nodC* sequences previously assigned to specific symbiovars (74). The alignment was then trimmed using trimAl version 1.4.rev22 (66) with the automated1 option. Finally, the trimmed alignment was used to construct a maximum likelihood phylogeny with IQ-TREE with the best-scoring substitution model, HKY+F+G4, identified by ModelFinder. Branch support was assessed using the Shimodaira–Hasegawa-like approximate likelihood ratio test (SH-aLRT) and ultrafast bootstrap analysis, with both metrics calculated from 1,000 replicates. Phylogenies were visualized using the iTOL webserver.

The NodC HMM pipeline was subsequently repeated using the full set of 216 *Rhizobium* genomes generated in this study, after which CD-HIT version 4.8.1 (75) was used to cluster the corresponding *nodC* sequences at a 99% sequence identity threshold to assign each isolate to a symbiovar subclade.

### Pangenome analysis

Panaroo version 1.5.0 (76) was used to calculate pangenomes for each of the 11 species. Panaroo was run in strict mode, with a sequence identity threshold of 90%, without edge filtering, and with the removal of invalid genes. The exception was *Rhizobium hidalgonense*, which was run in sensitive mode as strict mode failed due to there being only a single genome. Subsequently, the pangenomes for the 11 species were merged using the panaroo-merge function with a sequence identity threshold of 70%, a family sequence identity threshold of 50%, a length difference cut-off of 80%, and the merge paralogs option.

### Data availability

Raw sequencing data and annotated genome assemblies were deposited in the National Center for Biotechnology Information (NCBI) database under BioProject accession PRJNA1414687. Code to repeat all computational analyses described in this manuscript is available via GitHub at github.com/diCenzo-Lab/019_2026_Common_bean_rhizobia.

## RESULTS

### Ontario soils host diverse communities of rhizobia capable of nodulating common bean

Four cultivars of common bean were used in nodule trapping experiments to isolate common bean-nodulating rhizobia from nine soil samples collected across southern Ontario (**Table 1, Figure S1**). Depending on the soil and cultivar, plants inoculated with the soils ranged from appearing nitrogen deficient, with most nodules being smaller and white to slightly pink, to moderate or dark green with many larger and darker pink nodules (**Figures S2, S3**). These qualitative observations are consistent with most sites hosting many rhizobia capable of nodulating common bean, but with most rhizobia being low to moderate quality symbionts of this host species.

We recovered a total of 216 *Rhizobium* isolates, collecting between 21 and 26 isolates per soil and between 44 and 65 isolates per cultivar. Phylogenomic analyses indicated that these 216 isolates belong to 11 *Rhizobium* species, including two unnamed species (**Figure 1**), which formed 37 clusters when grouped at an ANI threshold of 99.8% (**Figure S4, Dataset S3**); increasing the ANI threshold to 99.9% did not split any of the 37 clusters. The number of isolates assigned to each species varied from 1 (*Rhizobium hidalgonense*) to 82 (*Rhizobium croatiense*) (**Table S1**). We note that since we preferentially isolated rhizobia from the “best” nodules per plant, and that there were biases in sampling sites (e.g., an over-representation of soils from the Huron Research Station), the taxonomic composition of our isolated rhizobia is unlikely to accurately represent the taxonomic composition of the broader population of common bean-nodulating rhizobia in Ontario.

**Figure 1.**
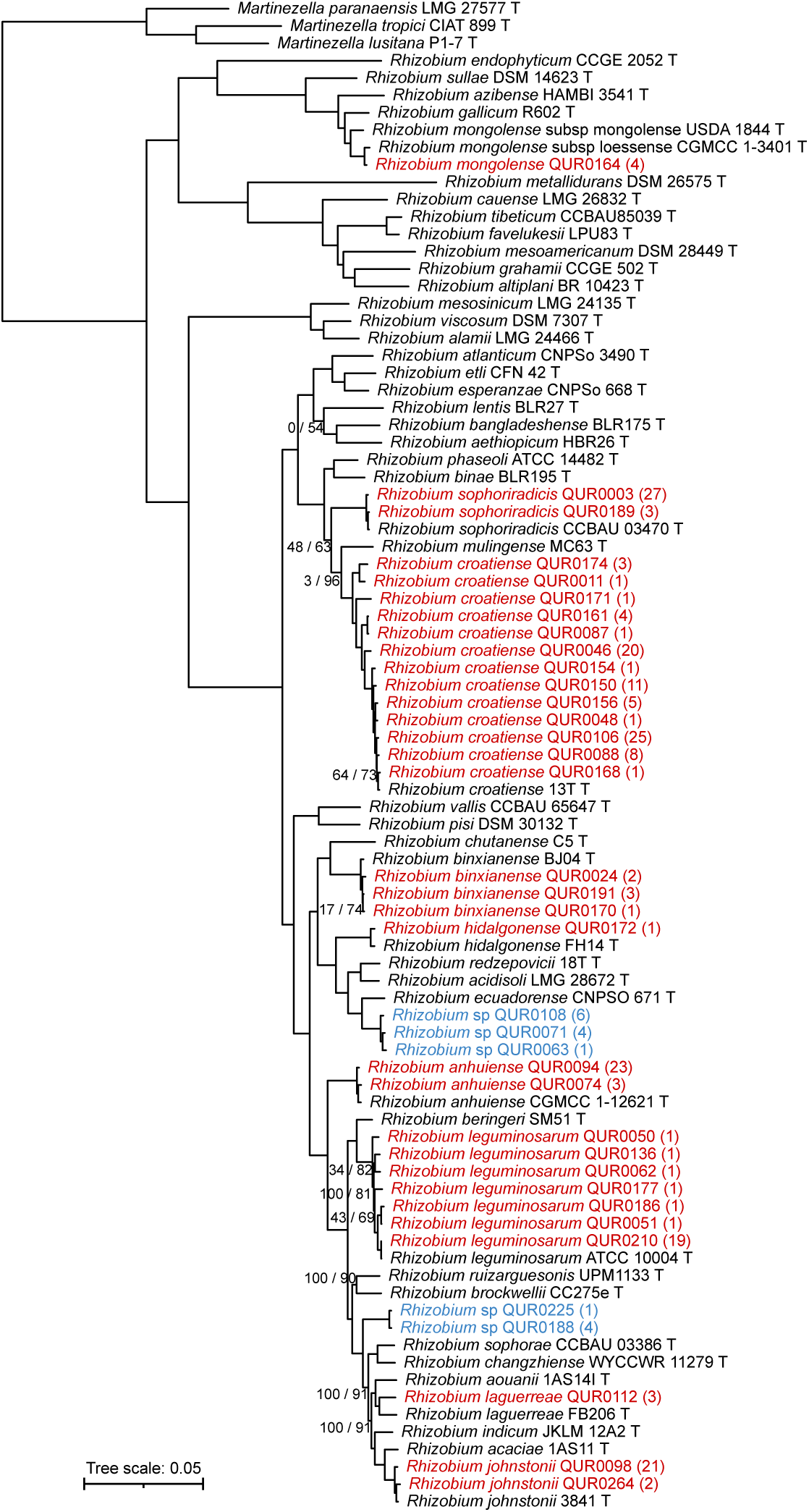
Phylogenomic analysis of the genus *Rhizobium*. A maximum likelihood core-proteome phylogeny of 37 representative *Rhizobium* isolates collected in this study together with all *Rhizobium* type strains with publicly available genome assemblies and rooted with type strains of the genus *Martinezella*. The 37 representative *Rhizobium* were selected based on dereplicating all 216 isolates at an ANI threshold of 99.8%; the number of isolates represented by each isolate is indicated in parentheses. Isolates collected in this study are shown in red (assigned to an existing species) or blue (assigned to an unnamed species). The numbers on the branches indicate values by SH-aLRT (left) and ultra-fast jackknife (right); values are only shown at nodes where at least one value is below 95%. The scale bar represents the average number of amino acid substitutions per site. An interactive version of this phylogeny is available at: itol.embl.de/shared/2L3P7mz1XsZy1.

Genome assemblies were generated for all 216 isolates, producing closed genomes for 136 (63%) isolates and high-quality draft genomes for all other isolates (**Dataset S1**). Based on the 136 closed genomes, our isolates carry between 4 and 8 replicons including the chromosome, and the number of replicons per genome varies both within and between species (between species Pr(>F) < 0.001; **Table S1** [Exp 1]) (**Table S2**). Genome size varies from ∼6.3 Mb to ∼7.8 Mb and differs significantly across species (Pr(>F) < 0.001; **Table S1** [Exp 1]), with *Rhizobium croatiense* having the smallest median genome size and *Rhizobium laguerreae* having the largest median genome size (**Table S2, Figure S5**). The pangenome of the 216 isolates includes a total of 20,528 gene groups, of which 4,281 (21%) belong to the core or soft-core genome (**Figure S6**). In addition, each of the 11 species contain between 48 and 1,926 unique gene groups present in their core or soft-core genome that are absent from the other 10 species (**Figure S7**).

### Symbiovar *phaesoli* is the dominant symbiovar across the *Rhizobium* isolates

Phylogenetic analysis of *nodC* revealed that nearly all (212 of 216) isolates belong to symbiovar *phaseoli* (**Figure 2A**). The exceptions are the four *R. mongolense* isolates, which belong to symbiovar *gallica* (previously known as symbiovar *gallicum*) (**Figure 2A**). Previous work defined multiple subclades of the symbiovar *phaseoli* based on *nodC* alleles (74, 77–79). Our isolates collectively carry three of these *nodC* alleles: 155 (72%) carry the γ-a allele, 29 (13%) carry the γ-b allele, and 28 (13%) carry the α allele (**Figure 2A**). In addition, examining the organization of *nod* genes in a subset of 23 representative isolates revealed five types of *nod* region organizations (**Figures 2B, S8**). Isolates of the symbiovar *phaseoli* have one of four types of *nod* region organization that correlate with an isolate’s symbiovar subclade classification, and these *nod* regions are most similar to Group C as defined by Eardly et al. (80). On the other hand, the *R. mongolense* isolate (symbiovar *gallica*) have a distinct *nod* region organization that is most similar to Group G as defined by Eardly et al. (80). Interestingly, *Rhizobium* species often include representatives belonging to multiple symbiovar *phaseoli* subclades; for example, *R. croatiense* and *R. leguminosarum* each have at least one representative from each of the three symbiovar *phaseoli* subclades (**Figures 2, S8, Table S3**). This observation suggests that there is extensive horizontal transfer of symbiotic genes between, and potentially within, common bean-nodulating *Rhizobium* species in Ontario.

**Figure 2.**
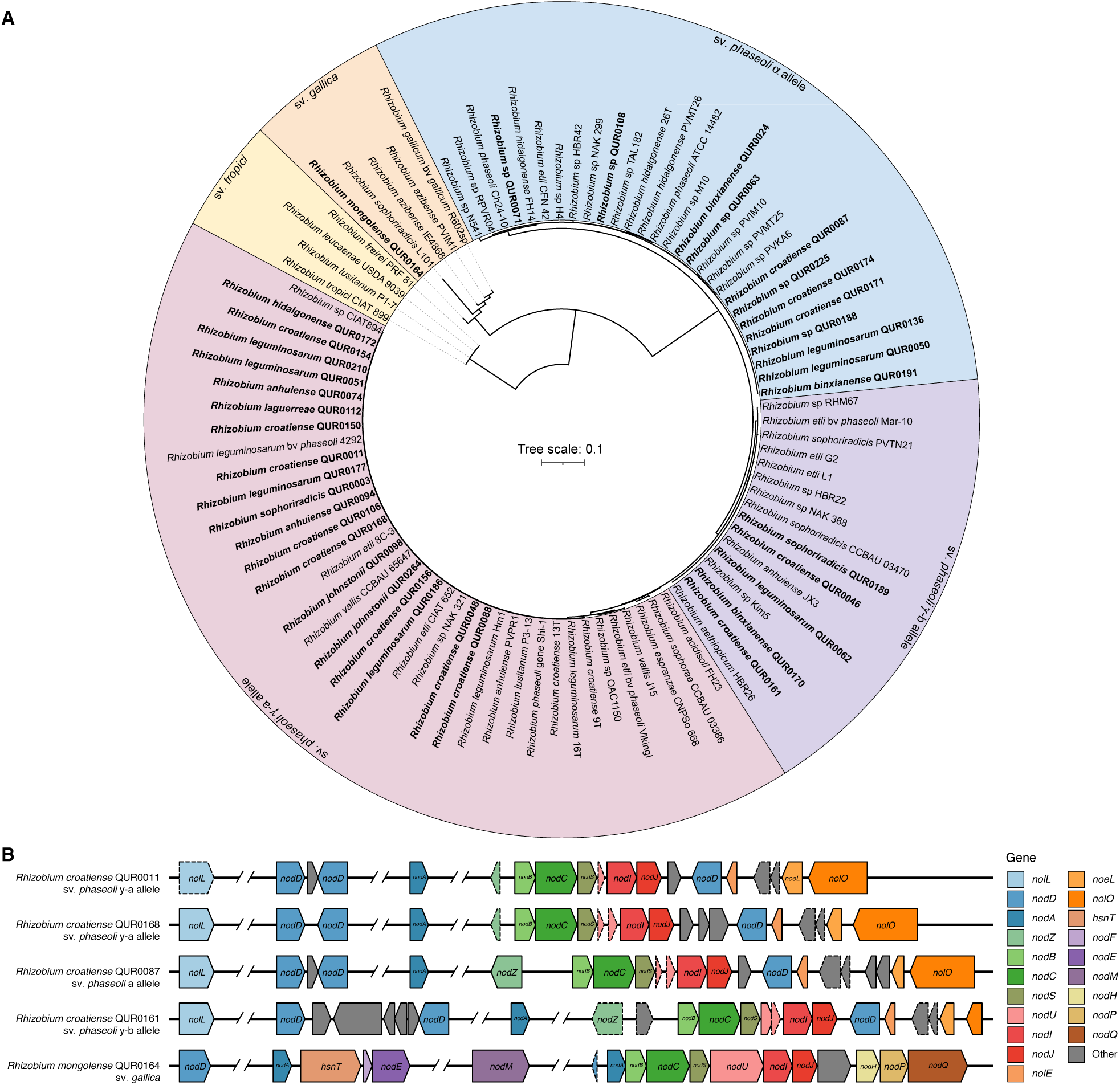
Organization and phylogenetic analysis of *nod* genes in representative *Rhizobium* isolates. (**A**) A maximum likelihood phylogeny of the *nodC* alleles of 37 representative *Rhizobium* isolates together with 54 *nodC* alleles from the literature. The scale bar represents the average number of nucleotide substitutions per site. An interactive version of this phylogeny, with support values, is available at: itol.embl.de/shared/2L3P7mz1XsZy1. (**B**) The organization of *nod* genes in five *Rhizobium* isolates representative of the five types of *nod* region organization observed (see **Figure S8** for additional examples). Genes are drawn to scale, orthologous genes are colour coded, and pseudogenes are indicated with dashed borders (note: *nolL* is not a pseudogene in all isolates with a *nod* gene organization represented by *Rhizobium croatiense* QUR0011). Gaps in sequences are represented by two angled lines with white space between them.

### Soil origin impacts the diversity of rhizobia recovered by nodule trapping

The taxonomic composition of the *Rhizobium* populations recovered by nodule trapping varied considerably across soil type (Pr(>F) = 0.001; **Table S1** [Exp 2]) (**Figure 3A**), with 70% of variation explained; none of the 11 rhizobial species were recovered from all nine soils. The rhizobial populations recovered from the five soils collected from southwestern Ontario, which also represent soils with a long history of common bean cultivation (**Table 1**), clustered together primarily due to a high relative abundance of *R. croatiense* (**Figures 3A, 3B**). Likewise, except for soil GSC149, the rhizobial populations recovered from the soils from south central and southeastern Ontario formed a cluster (**Figure 3A**). These rhizobial populations are unique in having a high relative abundance of *Rhizobium anhuiense* relative to other soils, while the rhizobial populations from south central Ontario uniquely have a high relative abundance of *Rhizobium sophoriradicis* (**Figure 3B**).

**Figure 3.**
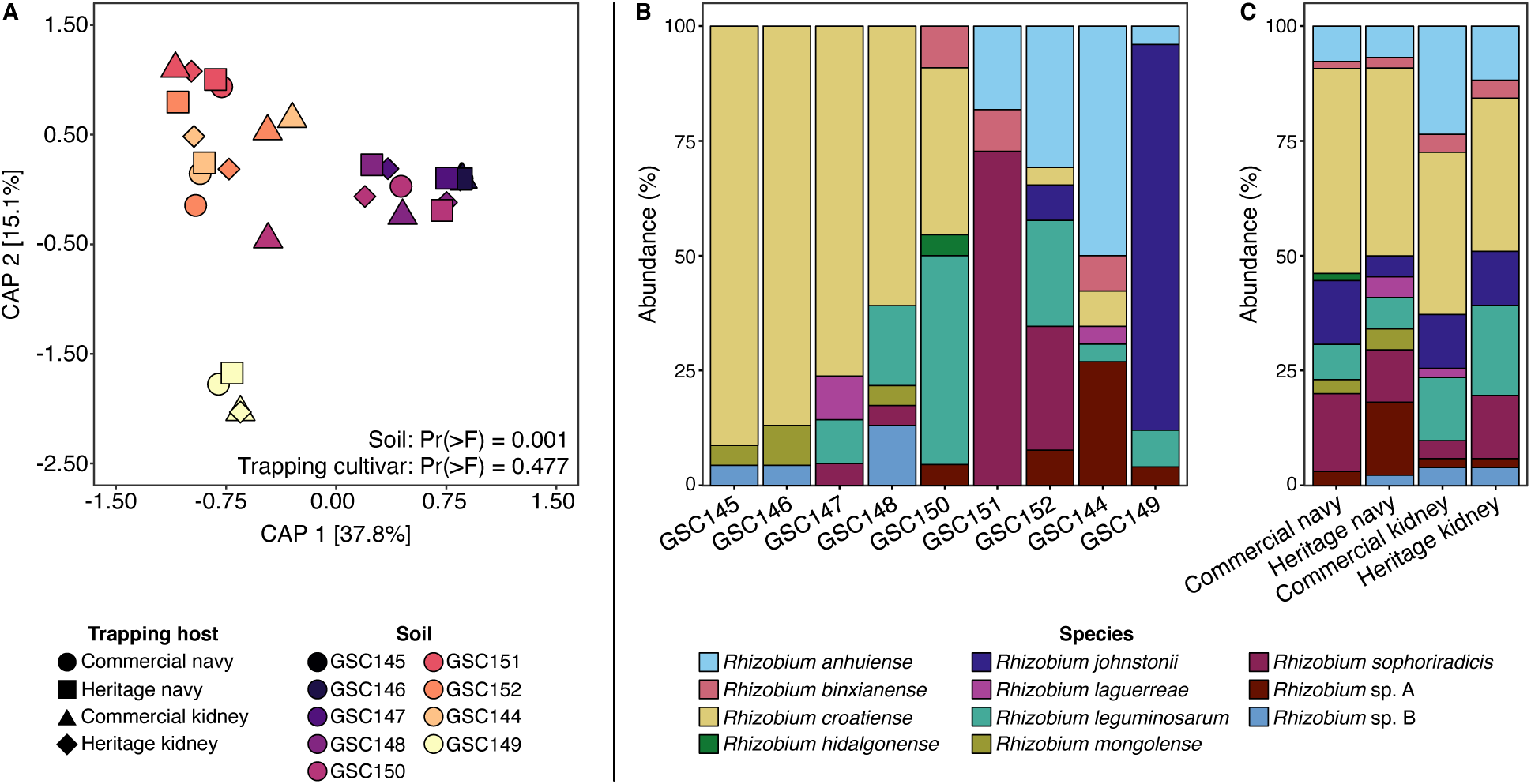
Impact of soil and common bean trapping cultivar on the diversity of the trapped rhizobia. (**A**) An ordination plot produced from a capscale analysis run on Bray-Curtis distances calculated based on the species-level composition of the rhizobial population recovered from each nodule trapping sample (N = 36). Shapes represent which common bean cultivar was used as the trapping host, while colours indicate which soil sample was used (**Table 1, Figure S1**). The p-values are from F-tests generated from running Type II ANOVAs on capscale analyses on models that included soil and trapping cultivar as main effects. (**B**, **C**) Stacked bar plots showing the composition of the recovered rhizobial populations based on (**B**) the original soil or (**C**) common bean cultivar used during the nodule trapping experiments.

On the other hand, there was no statistically significant effect of host trapping cultivar on the taxonomic composition of the recovered *Rhizobium* populations (Pr(>F) = 0.48, 3.2% of variation explained; **Table S1** [Exp 2]) (**Figures 3A, 3C**). However, it is worth noting that some biases in *Rhizobium* species recovery by host were observed. For example, *R. mongolense* (and thus symbiovar *gallica*) was only recovered when the navy bean cultivars were used for trapping; *R. leguminosarum* was recovered twice as frequently when using the kidney bean cultivars compared to the navy bean cultivars; and 64% of *Rhizobium* sp. A (all of which belong to symbiovar *phaseoli* subclade α) were isolated using the heritage navy bean cultivar for trapping (**Figure 3C**).

### Rhizobia from Ontario soils effectively fix nitrogen with common bean

The 216 *Rhizobium* isolates varied extensively in their ability to fix nitrogen in symbiosis with navy bean plants, with most isolates improving plant growth compared to nitrogen-free, uninoculated control plants (**Figure 4, Dataset S4**). Dried Shoot biomass significantly differed across both *Rhizobium* species and symbiovar sub-clade classifications (Pr(>F) < 0.001 and Pr(>F) = 0.01, respectively; **Table S1** [Exp 3]). The average quality of most species pairs represented in our collection were not statistically different from each other (**Figures 4, S9**). Two exceptions were *Rhizobium hidalgonense* (1 isolate) and *Rhizobium laguerreae* (3 isolates), which were statistically worse symbionts of common bean than most other species (**Figures 4, S9**). In addition, *Rhizobium* sp. A, whose isolates all belonged to the symbiovar *phaseoli* subclade α, were statistically better, on average, than all other species except for *R. mongolense* and *Rhizobium johnstonii* (**Figures 4, S9**); the average shoot dry weight of plants inoculated with *Rhizobium* sp. A was 2.3 g/plant (median of 2.5 g/plant), which is ∼38% larger than the overall average of all inoculated plants of 1.7 g/plant (median of 1.6 g/plant). Note, however, that although the average effectiveness of isolates of this unnamed species was higher than the overall average, the top performing isolates from this species did not outperform the top isolates from the other *Rhizobium* species (**Figure 4A**). In addition, isolates of the symbiovar *phaseoli* subclade γ-a (which was the most abundant subclade in our dataset) were, on average, worse partners of common bean than isolates of the symbiovar *phaseoli* subclades γ -b and α (**Figures 4, S10**).

**Figure 4.**
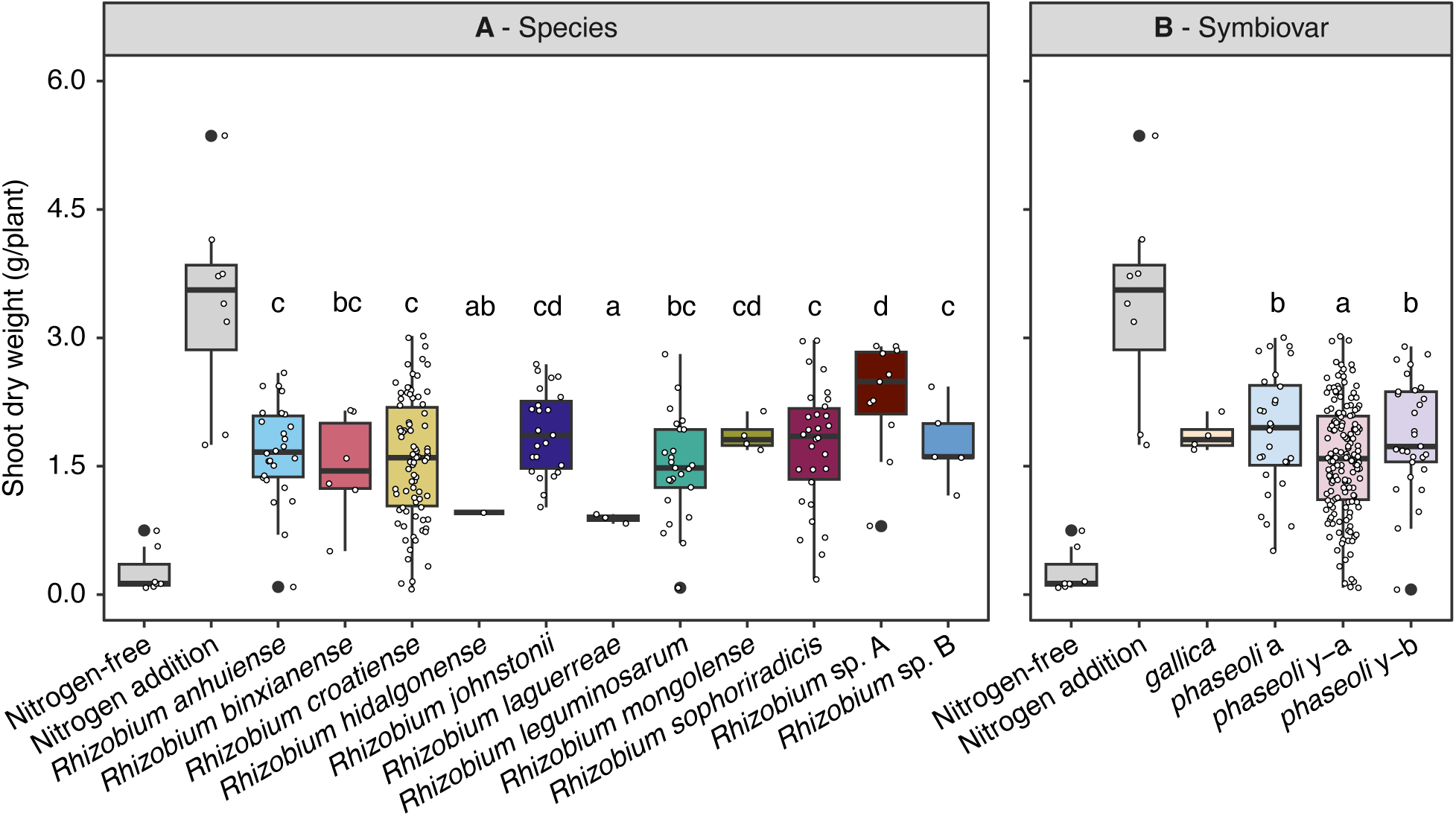
Symbiotic efficiency of *Rhizobium* isolates from southern Ontario. Variation in shoot dry weights of navy bean plants inoculated with one of 216 *Rhizobium* isolates, summarized either by (**A**) species or (**B**) symbiovar subclade. In both panels, each dot represents a single plant inoculated with one of the 216 *Rhizobium* isolates, while the boxplots represent the distribution of the data for a given species or symbiovar subclade. The “nitrogen-free” and “nitrogen addition” conditions represent uninoculated plants either without or with nitrogen supplementation, respectively. Letters represent statistically significant groupings based on pairwise comparison of the estimated marginal means followed by a Tukey’s multiple test correction (p-value < 0.05); the control treatments were excluded from the statistical analysis.

In addition to measuring shoot dry weight at harvest (5 weeks post inoculation), weekly NDVI measurements of each plant were taken. Week 3 was the first week where the negative controls were visibly worse (more chlorotic) than inoculated plants, and so we focused our analysis of the NDVI values on the week 3 data. Week 3 NDVI was a statistically significant predictor of final plant shoot dry weights (Pr(>F) < 0.001; **Table S1**). Indeed, the mean shoot dry weight of plants in the bottom quartile of week 3 NDVI values was 1.01 (median 1.00) compared to a mean shoot dry weight of 1.95 (median 1.92) for plants in the top quartile of week 3 NDVI values (**Figure S11**). Overall, this trend suggests that early nodule formation and nitrogen fixation was associated with higher cumulative nitrogen fixed over the five-week experiment.

To validate the general patterns observed in the initial screen of the nitrogen fixation capacity of the 216 *Rhizobium* isolates, a follow-up experiment was performed with 12 isolates: eight (from two species) that were expected to be good quality partners, two (from one species) that were expected to be medium quality partners, and two (from one species) that were expected to be poor quality partners, all based on the results from the first greenhouse experiment. A starter of 5 mM of nitrogen was included in these experiments to boost early plant health (see next section for rationale). There was a statistically significant effect of *Rhizobium* strain on the shoot dry weight of the inoculated plants (Pr(>F) < 0.001; **Table S1** [Exp 4]) (**Figure S12**). Most results were consistent with trends observed in the initial screen; the two *Rhizobium laguerreae* isolates were poor quality symbionts with navy bean, while the four *R. croatiense* isolates and the four isolates belonging to unnamed *Rhizobium* sp. A were generally high quality symbionts (**Figure 5, Dataset S5**). However, the two *R. mongolense* isolates that were expected to be medium quality symbionts were not statistically different from the *R. croatiense* and *R. sp*. isolates (**Figures 5, S12**). Nevertheless, the overall alignment of these results from those of the initial screen demonstrate that the *Rhizobium* collected in this study vary in partner quality, with many isolates being good quality partners of navy bean plants.

**Figure 5.**
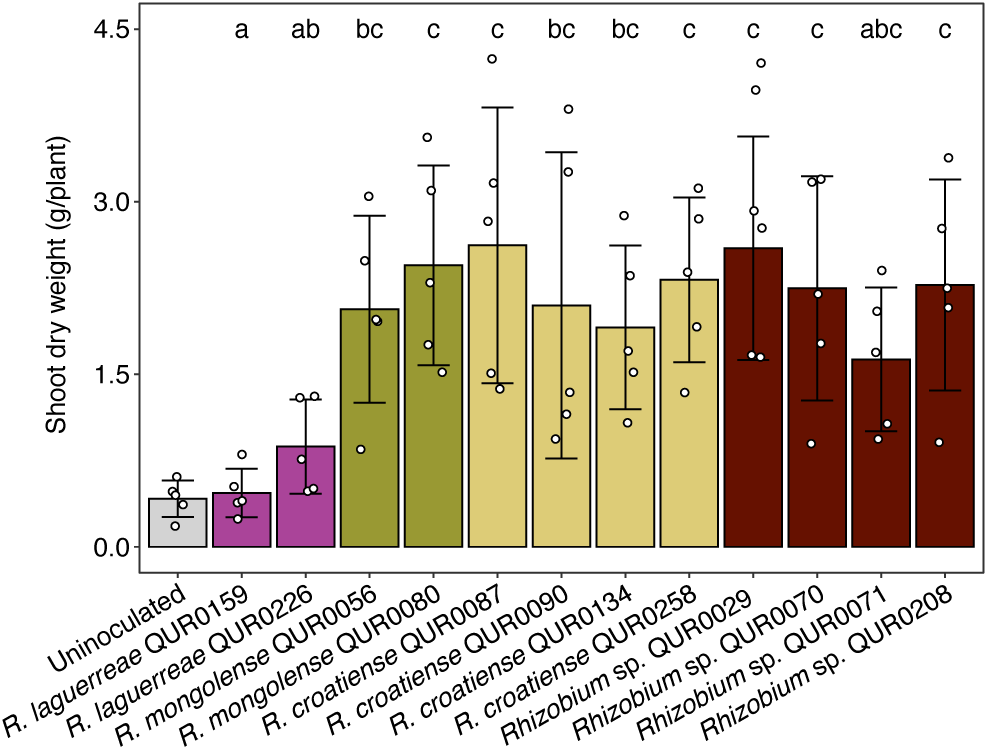
Symbiotic efficiency of a subset of *Rhizobium* isolates. Variation in shoot dry weights of navy bean plants inoculated with one of 12 *Rhizobium* isolates. Each dot represents a single plant inoculated with a given *Rhizobium* isolate, while the columns and error bars indicate the mean and standard deviation, respectively. The “uninoculated” condition represents plants without *Rhizobium* inoculation. Letters represent statistically significant groupings based on pairwise comparison of the estimated marginal means followed by a Tukey’s multiple test correction (p-value < 0.05); the uninoculated controls were excluded from the statistical analysis.

### *Rhizobium* inoculation synergizes with limited nitrogen addition in the lab

Given the delay between inoculation of plants and the onset of nitrogen fixation, we examined whether supplementing pots with nitrogen would improve plant health and synergize with *Rhizobium* inoculation. Two nitrogen regiments, each with variation in the concentration of supplemental nitrogen, were tested: ongoing nitrogen fertilization and one-time fertilization at the start of the experiment.

In the ongoing nitrogen fertilizer regimen, plant shoot dry weight was significantly impacted by nitrogen concentration (Pr(>F) = 0.005), rhizobium inoculation (Pr(>F) < 0.001), and their interaction (Pr(>F) < 0.001) (**Table S1** [Exp 5a]). There was a beneficial effect of combining nitrogen addition and rhizobium inoculation at low nitrogen concentrations (**Figure 6A**); however, increasing nitrogen concentrations above 1 mM resulted in decreasing benefit of inoculation (measured as a gain in biomass relative to uninoculated plants experiencing the same nitrogen treatment), with no benefit of inoculation observed at 10 mM or 20 mM nitrogen (**Figure 6B**).

**Figure 6.**
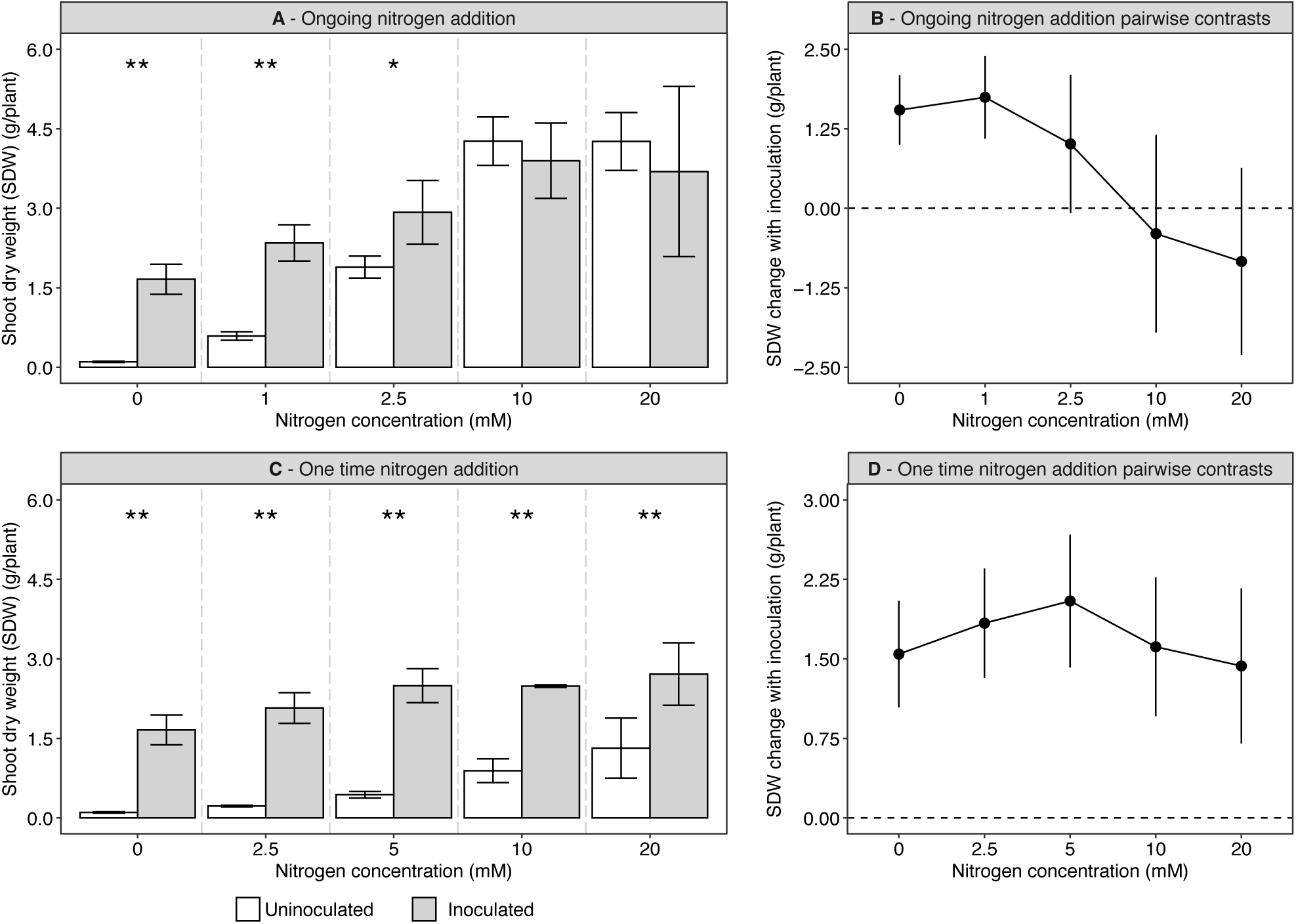
Impact of nitrogen supplementation on the benefit of *Rhizobium* inoculation. (**A**) Navy bean plants were given various concentrations of nitrogen supplementation on an (**A**) ongoing basis or (**C**) one time at the start of the experiment, and the plants were either inoculated (grey) or not (white) with *Rhizobium* sp. QUR0071. Columns and error bars represent the mean and standard deviation of triplicate plants, respectively. Asterisks indicate the significance level of pairwise contrasts between the inoculated and uninoculated control plants at a given concentration of nitrogen: *, Pr(>F) < 0.1; ** Pr(>F) < 0.001). Pairwise contrasts between the inoculated and uninoculated plants, with their 95% confidence intervals, are shown for the plants receiving nitrogen supplementation on an (**B**) ongoing basis or (**D**) one time at the start of the experiment. Values above 0 indicate that the inoculated plants were larger than the uninoculated plants, whereas values below 0 indicated the uninoculated plants were larger than the inoculated plants.

On the other hand, for the one-time nitrogen fertilizer regimen, plant shoot dry weight was significantly impacted by *Rhizobium* inoculation (Pr(>F) < 0.001) and the interaction between rhizobium inoculation and nitrogen concentration (Pr(>F) < 0.001), but not nitrogen alone (Pr(>F) = 0.12) (**Table S1** [Exp 5b]). In contrast to the ongoing nitrogen fertilizer regimen, there was a beneficial effect of rhizobium inoculation at all nitrogen concentrations (**Figure 6C**). Notably, there was a positive correlation between the concentration of the added nitrogen and the benefit of inoculation up to 5 mM of nitrogen (**Figure 6D**), suggesting that the addition of small amounts of nitrogen at the start of the experiment facilitated more rapid establishment of nitrogen-fixing nodules.

## DISCUSSION

To identify effective rhizobial isolates for development into commercial inoculants for common bean in Canada, we characterized 216 rhizobia originating from southern Ontario soils. We conducted whole genome comparisons across the isolates to taxonomically classify them and assign them to symbiovars, based on the *nodC* locus. We also screened isolates based on their performance as symbionts when paired with a commercial navy bean variety in greenhouse experiments. We uncovered three main results. First, there was substantial species diversity among *Rhizobium* strains in southern Ontario that can form nodules on common bean. Second, the composition of *Rhizobium* species associated with common bean varied across soil samples. Third, we identified an unnamed *Rhizobium* species whose members are on average better common bean symbionts than isolates from other *Rhizobium* species. Our results have important implications for the development of a successful inoculum for use with common bean in Ontario agriculture, which we discuss below.

### Southern Ontario soils host a diversity of *Rhizobium* symbionts of common bean

Common bean is often referred to as a promiscuous host as it is well known to associate with a diversity of *Rhizobium* species (12). Consistent with common bean having a broad symbiont breadth, we recovered 11 distinct species from our nodule trapping study, including two unnamed species. The recovery of a high diversity of species is in line with other studies showing that soils in many jurisdictions – including in Europe (Greece, Croatia), Africa (Ethiopia, Uganda), Central and South America (Mexico, Ecuador, Brazil), and Asia (China) – host a diversity of rhizobia able to nodulate common bean (74, 81–86).

Although common bean is not native to Canada, it is estimated to have been introduced to southern Ontario from Mexico by Indigenous communities prior to CE 1250 (87, 88). The initial introduction of *P. vulgaris* to Ontario may have resulted in the co-introduction of one or more compatible *Rhizobium* strains, from which the required symbiotic genes could have spread horizontally to native rhizobia, thereby creating an increasingly diverse suite of rhizobia able to nodulate common bean (89–92). Our data are consistent with this scenario. There are at least seven reported *Rhizobium* symbiovars capable of nodulating common bean (93). Of these, symbiovar *phaseoli* dominates in the centres of diversification, with *nodC* alleles α and γ dominating in Mesoamerica and *nodC* allele δ dominating in the Andes and Ecuador (77). In our study, 98% (212 of 216) isolates carried the *nodC* alleles α, γ-a, or γ-b, consistent with the symbiotic genes carried by these isolates originating in Mesoamerica and subsequently being introduced to Ontario with the spread of Indigenous Three Sisters agricultural practices. However, common bean-nodulating rhizobia in Mesoamerica were classically thought to be dominated by the species *Rhizobium etli* (77), whereas no *R. etli* isolates were recovered in our nodule trapping experiments. While we cannot rule out that the lack of *R. etli* could be due to the specific cultivars we used for trapping, we instead interpret our results as being partially driven by competition between introduce rhizobia that travelled with common bean versus native rhizobia populations, and that the common bean rhizobia observed in southern Ontario currently are the products of horizontal spread of symbiosis genes from the introduced (and subsequently lost) rhizobia to native rhizobial populations. However, we note that many isolates previously classified as *R. etli* have been renamed (85, 94), and thus, we cannot rule out that some or all our *Rhizobium sophoriradicis* and *R. croatiense* isolates (which fall within the broader *R. etli* clade) are descendants of introduced rhizobia.

### Biogeographical structuring of common bean-nodulating *Rhizobium* across southern Ontario

We preferentially isolated rhizobia from large, pink nodules during our nodule trapping experiments. While this strategy allowed us to enrich our collection in effective nitrogen-fixing rhizobia, it also means that the recovered populations are likely not representative of the typical taxonomic composition of common bean nodulating rhizobia in each soil. Nevertheless, with this caveat in mind, we found that soil sample, rather than common bean cultivar, was a stronger predictor of the species-level composition of *Rhizobium* populations recovered from *P. vulgaris* nodules. Likewise, previous studies have shown genetic differentiation among common bean-associated rhizobia from different geographic regions (95–97). The variation in rhizobia recovered from each soil could reflect variation in soil or climatic characteristics, such as soil pH, leading to enrichment of species adapted to the local environment at each sampling site (98, 99). Indeed, we found that there was a spatial grouping of sites, with the rhizobial populations recovered from southwestern Ontario soil samples grouping together and separate from samples collected in southcentral or southeastern Ontario. However, this spatial pattern was confounded by common bean cultivation history, as the southwestern Ontario sites (the Huron Research Station and a commercial common bean farm) were the only sites known to have a recent, long history of common bean cultivation. Thus, we cannot differentiate whether the similarity in the recovered rhizobial populations from these sites was due to shared soil or environmental factors, or due to a shared history of common bean cultivation enriching specific rhizobial species.

Other studies have demonstrated that host genotype can influence symbiont specificity in *P. vulagris*. For instance, a local bean cultivar was found to recruit a more diverse set of rhizobia from its local soil compared to a foreign cultivar (96). Similarly, beans from the Mesoamerican and Andean gene pools differed in their specificity for *Rhizobium etli* strains based on which *nodC* allele they possessed (95). In contrast, despite using navy bean and kidney bean plants for our nodule trapping experiments, which belong to the Mesoamerican and Andean gene pools, respectively (100), no statistically significant impact of host cultivar on rhizobial species composition was detected. The lack of a host effect could be due to a lack of statistical power, since we only included one replicate for each soil – cultivar pairing. Alternatively, we note that both studies cited above were conducted within the centers of diversity for common bean, where long term coevolution between local hosts and rhizobia could enhance host genotype effects. In contrast, neither navy bean nor kidney bean is native to our soil sampling sites.

### Within and between *Rhizobium* species variation in nitrogen-fixing capacity with common bean

There was substantial variation in the partner quality of our *Rhizobium* isolates, both within and between species. While the average partner quality of most of the recovered *Rhizobium* species did not statistically differ, the initial screen of partner quality suggested that isolates from one of the novel *Rhizobium* are on average better symbionts of navy bean plants than the rest of our collection. In future work, it will be interesting to understand the genetic basis for the elevated nitrogen-fixing capacity of this species with common bean, and to explore whether its beneficial effect holds true with other common bean cultivars or under field conditions.

More broadly, most of our isolates were effective nitrogen fixers with navy bean plants. However, our isolation procedure meant that we selectively enriched for effective partners and thus we cannot comment on whether this is true more broadly of the common bean-nodulating rhizobial population in Ontario. In fact, most nodules visually appeared to be poorly effective in our initial nodule trapping experiments, suggesting that most common bean-nodulating rhizobia in Ontario soils are poor-quality symbionts of this legume. This result is not be surprising given it is known that common bean often associates with rhizobia that are poor nitrogen fixers on this host (12). In addition, the majority of our sampling sites had limited recent history of common bean cultivation, and thus, there may have been a lack of selective pressure to increase the abundance of effective rhizobia in these soils (101, 102).

### Towards an inoculant for Ontario common bean crops

The primary goal of this study was to establish a genome-sequenced collection of Ontario common bean-nodulating rhizobia to support future studies aiming to identify isolates with the potential for commercialization as novel inoculants, thereby reducing reliance on nitrogen fertilization of common bean crops in Ontario. The identification of isolates with high nitrogen-fixation capacity with navy bean plants is a good first step towards the development of new commercial products. However, various potential hurdles remain, necessitating further studies.

To date, we have only screened our *Rhizobium* library on a single navy bean under controlled greenhouse conditions. Previous studies have detected significant *Rhizobium* genotype by common bean genotype (GxG) interactions (103, 104), suggesting that the isolates that performed well with navy bean plants may not be top performers with other cultivars. In addition, the taxonomic differences in the *Rhizobium* communities recovered from each soil suggests the potential for *Rhizobium* genotype by environment (GxE) interactions or *Rhizobium* genotype by common bean genotype by environment (GxGxE) interactions (105, 106), which could reduce how reliably a given strain performs across various agro-climatic conditions. As our initial experiments were performed in a vermiculite – sand potting mixture in a greenhouse, they do not reflect realistic field conditions. As such, the isolates that performed best in our initial screen may not be the top performers in other conditions, and the top performers may vary in different soils and/or with different cultivars. Consequently, screening for isolates that reliably perform well across cultivars and soil types, under varying environmental conditions, will be an important next step towards identifying which isolates are best candidates for commercialization.

Beyond nitrogen fixation effectiveness, it is also important to consider the competitive ability of each isolate. The experiments reported here all involved single strain inoculation experiments, which precludes having to compete for access to nodules. The competitiveness, or lack thereof, of commercial inoculants for nodule occupancy has long been recognized as a potential hurdle to their success under field conditions (29, 31, 107). Rhizobia can independently vary in their partner quality (amount of nitrogen fixed) and their competitiveness for nodule occupancy traits. Indeed, partner quality and competitiveness were uncorrelated in a set of common bean-nodulating rhizobia from Kenya (108). Our nodule trapping experiments suggest that Ontario soils host a resident population of common bean-nodulating rhizobia, and that many of these strains are likely to be poor-quality symbionts. Therefore, any new inoculant would need to be capable of outcompeting these suboptimal resident rhizobia for nodule occupancy. Consequently, co-inoculation experiments will need to be conducted to test whether these high-quality partners are also highly competitive against isolates (of varying partner quality) naturally present in non-sterile Ontario soils.

## ACKNOWLEDGEMENTS

We thank Hensall Co-op and the Kingston Area Seed System Initiative (KASSI) for kindly providing the commercial and heritage common bean varieties, respectively. This research was also supported by the “Bio-inoculants for the promotion of nutrient use efficiency and crop resiliency in Canadian agriculture” project funded by the Government of Canada through Genome Canada and Genome Prairie (CSAFS-ICT 19308), the Government of Ontario through an Ontario Research Fund - Interdisciplinary Challenge Teams (ORF-ICT) – Climate Action Genomics Initiative grant (File ICT 19308), and the Ontario Bean Growers.

