## Supplementary Materials for "Isolation of rhizobia from Ontario soils that are effective at fixing nitrogen with common bean (*Phaseolus vulgaris*)"

**This PDF file includes:**

Tables S1-S3 (pages 2-4)
Figures S1-S12 (pages 5-16)
Legends for Datasets S1-S6 (page 17)

**Other supplementary materials for this manuscript include the following:**

Datasets S1-S4 (Excel file)

**Table S1.** Results of all (mixed) linear models and capscale analyses run as part of this study.

| **Exp. 1 - Genome size and replicon counts** | | | | | |
|---|---|---|---|---|---|
| Model | Response | Predictor | d.f. | F value | Pr(>F) |
| Model 1 | Genome size | *Rhizobium* species | 10 | 156.64 | < 2.2e-16 |
| Model 2 | Replicon count | *Rhizobium* species | 9 | 57.028 | < 2.2e-16 |
| **Exp. 2 - Nodule trapping (capscale)** | | | | | |
| Model | Response | Predictor | d.f. | F value | Pr(>F) |
| Model 3 | - | Source soil | 8 | 7.9921 | 0.001 |
| Model 3 | - | Trapping host | 3 | 0.969 | 0.454 |
| **Exp. 3 - Initial screen of *Rhizobium* isolates on common bean** | | | | | |
| Model | Response | Predictor | d.f. | F value | Pr(>F) |
| Model 4 | Shoot dry weight | *Rhizobium* species | 10 | 11.944 | 3.643e-16 |
| Model 5 | Shoot dry weight | *Rhizobium* symbiovar | 2 | 4.6424 | 0.01065 |
| Model 6 | Shoot dry weight | Week 3 NDVI | 1 | 144.67 | < 2.2e-16 |
| **Exp. 4 - Follow-up screen of *Rhizobium* isolates on common bean** | | | | | |
| Model | Response | Predictor | d.f. | Chisq | Pr(>Chisq) |
| Model 7 | Shoot dry weight | Strain (fixed) | 11 | 70.483 | 9.887e-11 |
| Model 7 | Shoot dry weight | Block (random) | 1 | 32.073 | 1.485e-08 |
| **Exp. 5a - Effect of ongoing nitrogen supplementation** | | | | | |
| Model | Response | Predictor | d.f. | F value | Pr(>F) |
| Model 8 | Shoot dry weight | Nitrogen concentration | 4 | 5.423 | 0.004801 |
| Model 8 | Shoot dry weight | Inoculation (yes/no) | 1 | 99.439 | 9.326e-09 |
| Model 8 | Shoot dry weight | Interaction term | 4 | 40.664 | 8.810e-09 |
| **Exp. 5b - Effect of one-time nitrogen supplementation** | | | | | |
| Model | Response | Predictor | d.f. | F value | Pr(>F) |
| Model 9 | Shoot dry weight | Nitrogen concentration | 4 | 2.1173 | 0.1184 |
| Model 9 | Shoot dry weight | Inoculation (yes/no) | 1 | 493.7362 | 4.660e-15 |
| Model 9 | Shoot dry weight | Interaction term | 4 | 21.9595 | 6.611e-07 |

**Table S2.** Species-level summary of the genome size and replicon count for the *Rhizobium* isolates collected in this study.

| Species | No. of isolates | Genome size (Mb) | | | No. of replicons * | | |
|---|---|---|---|---|---|---|---|
| | | Minimum | Median | Maximum | Minimum | Median | Maximum |
| *R. anhuiense* | 26 | 7.04 | 7.30 | 7.57 | 4 | 5 | 7 |
| *R. binxianense* | 6 | 6.86 | 6.98 | 7.01 | ND † | ND | ND |
| *R. croatiense* | 82 | 6.30 | 6.63 | 7.23 | 6 | 7 | 8 |
| *R. hidalgonense* | 1 | 6.87 | 6.87 | 6.87 | 4 | 4 | 4 |
| *R. johnstonii* | 23 | 7.45 | 7.59 | 7.61 | 6 | 6 | 7 |
| *R. laguerreae* | 3 | 7.59 | 7.60 | 7.60 | 7 | 7 | 7 |
| *R. leguminosarum* | 25 | 7.41 | 7.58 | 7.84 | 5 | 5 | 6 |
| *R. mongolense* | 4 | 7.26 | 7.26 | 7.33 | 4 | 4 | 4 |
| *R. sophoriradicis* | 30 | 6.89 | 6.90 | 7.40 | 6 | 6 | 6 |
| *Rhizobium* sp. A | 11 | 6.81 | 7.21 | 7.21 | 7 | 8 | 8 |
| *Rhizobium* sp. B | 5 | 7.31 | 7.36 | 7.52 | 6 | 6 | 6 |

* Calculated only based on complete assemblies (i.e., excluding draft assemblies).

† ND – not determined as no genomes within this species were marked as complete.

**Table S3.** Distribution of symbiovars across the 11 *Rhizobium* species collected in this study.

| Species | Symbiovar | | | |
|---|---|---|---|---|
| | *phaseoli* γ-a | *phaseoli* γ-b | *phaseoli* α | *gallica* |
| *Rhizobium anhuiense* | 26 | 0 | 0 | 0 |
| *Rhizobium binxianense* | 0 | 1 | 5 | 0 |
| *Rhizobium croatiense* | 53 | 24 | 5 | 0 |
| *Rhizobium hidalgonense* | 1 | 0 | 0 | 0 |
| *Rhizobium johnstonii* | 23 | 0 | 0 | 0 |
| *Rhizobium laguerreae* | 3 | 0 | 0 | 0 |
| *Rhizobium leguminosarum* | 22 | 1 | 2 | 0 |
| *Rhizobium mongolense* | 0 | 0 | 0 | 4 |
| *Rhizobium sophoriradicis* | 27 | 3 | 0 | 0 |
| *Rhizobium* sp. A | 0 | 0 | 11 | 0 |
| *Rhizobium* sp. B | 0 | 0 | 5 | 0 |

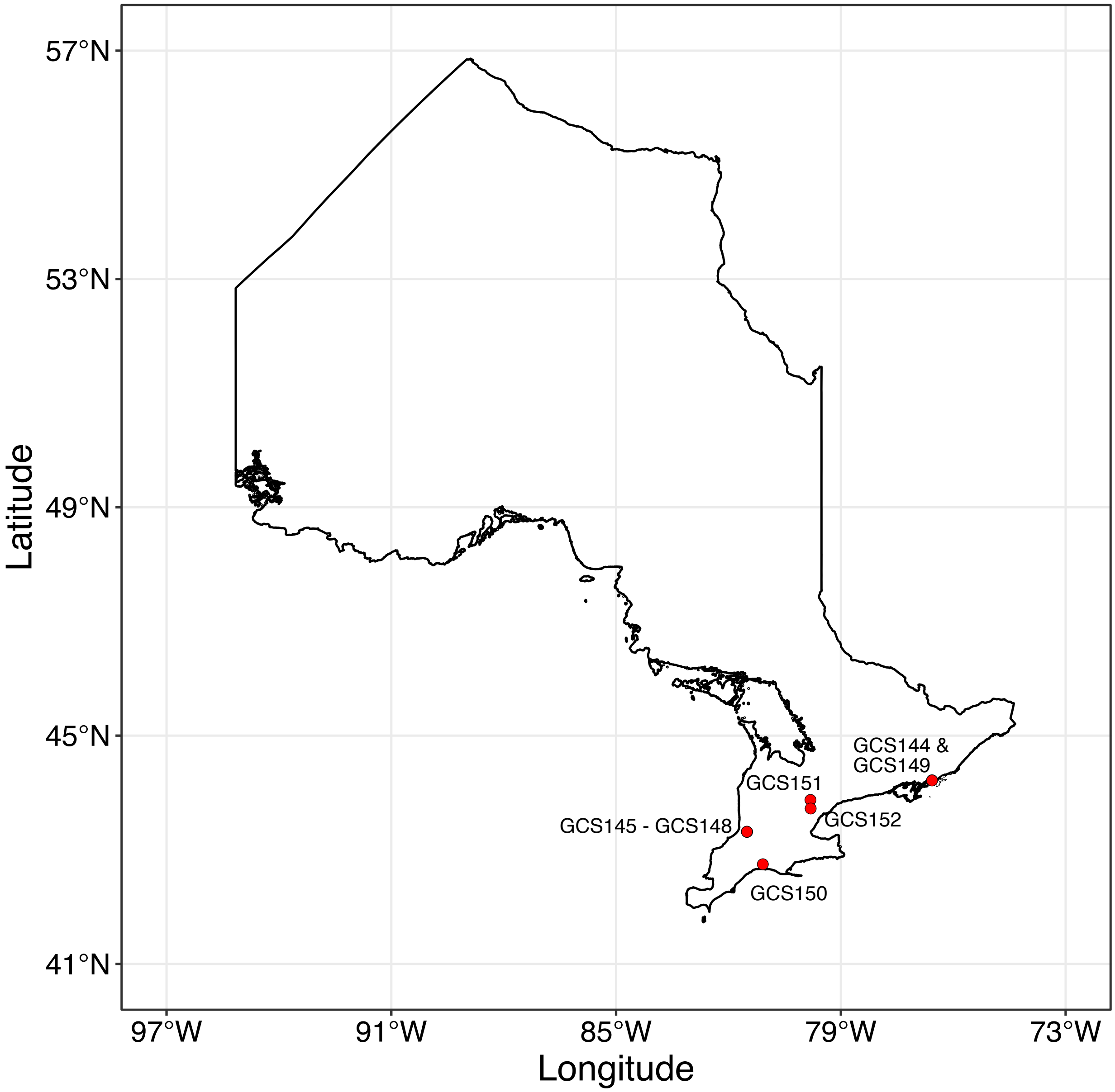


**Figure S1. Soil sampling sites.** A map of Ontario, Canada, is provided showing the locations where the nine soil samples were collected (red dots). The dots for the soil samples GCS145-148 overlap, as do GCS144 and GCS149, and thus individual dots are not apparent for these samples.

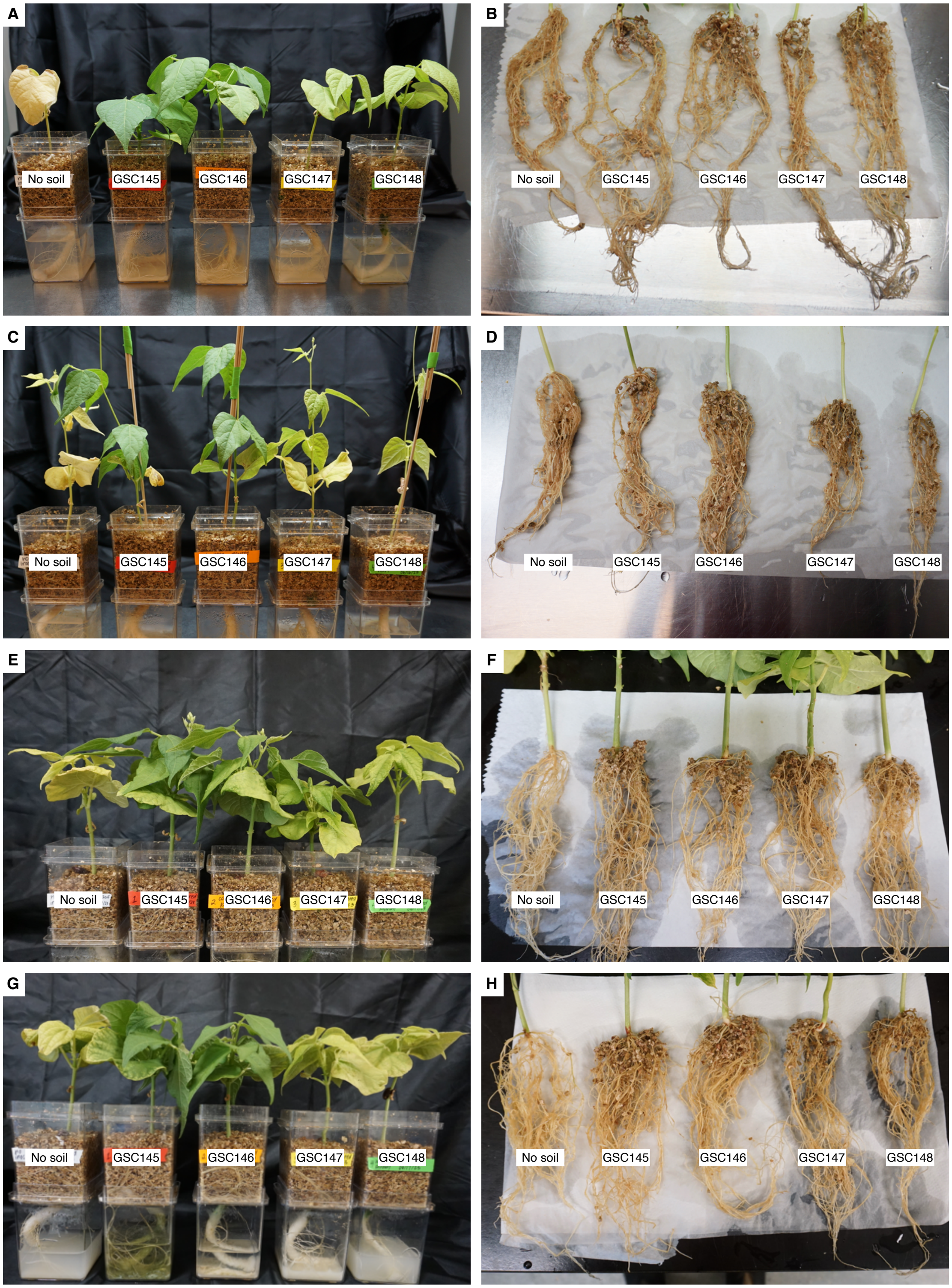


**Figure S2. Nodule trapping from soils collected at the Huron Research Station.** Photos of plants four weeks post inoculation with soils are shown. Photos are given for (**A**,**B**) commercial navy bean (AAC Shock), (**C**,**D**) heritage navy bean, (**E**,**F**) commercial kidney bean (Dynasty), and (**G**,**H**) heritage kidney bean. Plants are labelled based on the soil added to the pots, and the soils are described in **Table 1** and **Figure S1**.

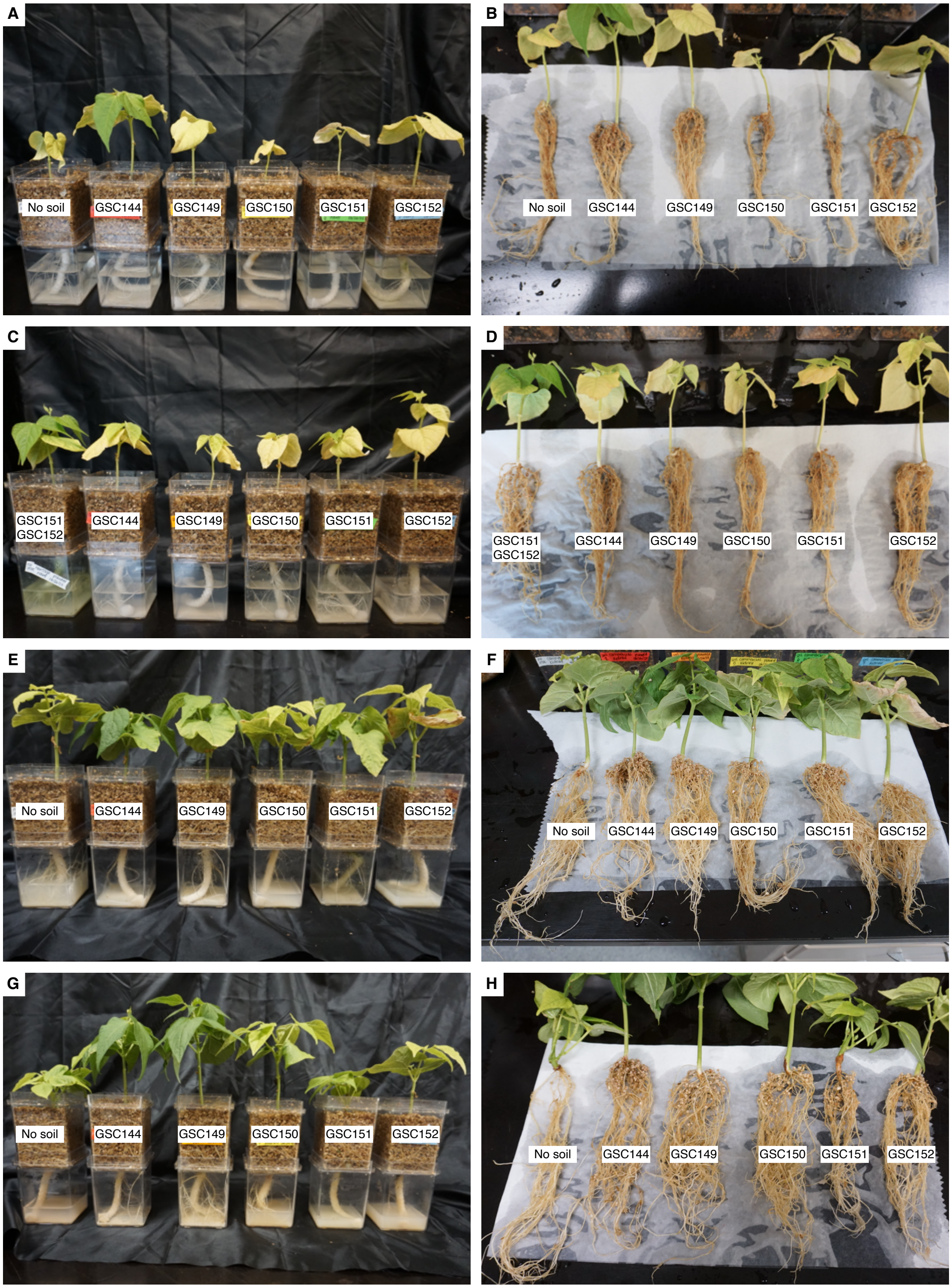


**Figure S3. Nodule trapping from soils collected other southern Ontario soils.** Photos of plants four weeks post inoculation with soils are shown. Photos are given for (**A**,**B**) commercial navy bean (AAC Shock), (**C**,**D**) heritage navy bean, (**E**,**F**) commercial kidney bean (Dynasty), and (**G**,**H**) heritage kidney bean. Plants are labelled based on the soil added to the pots, and the soils are described in **Table 1** and **Figure S1**.

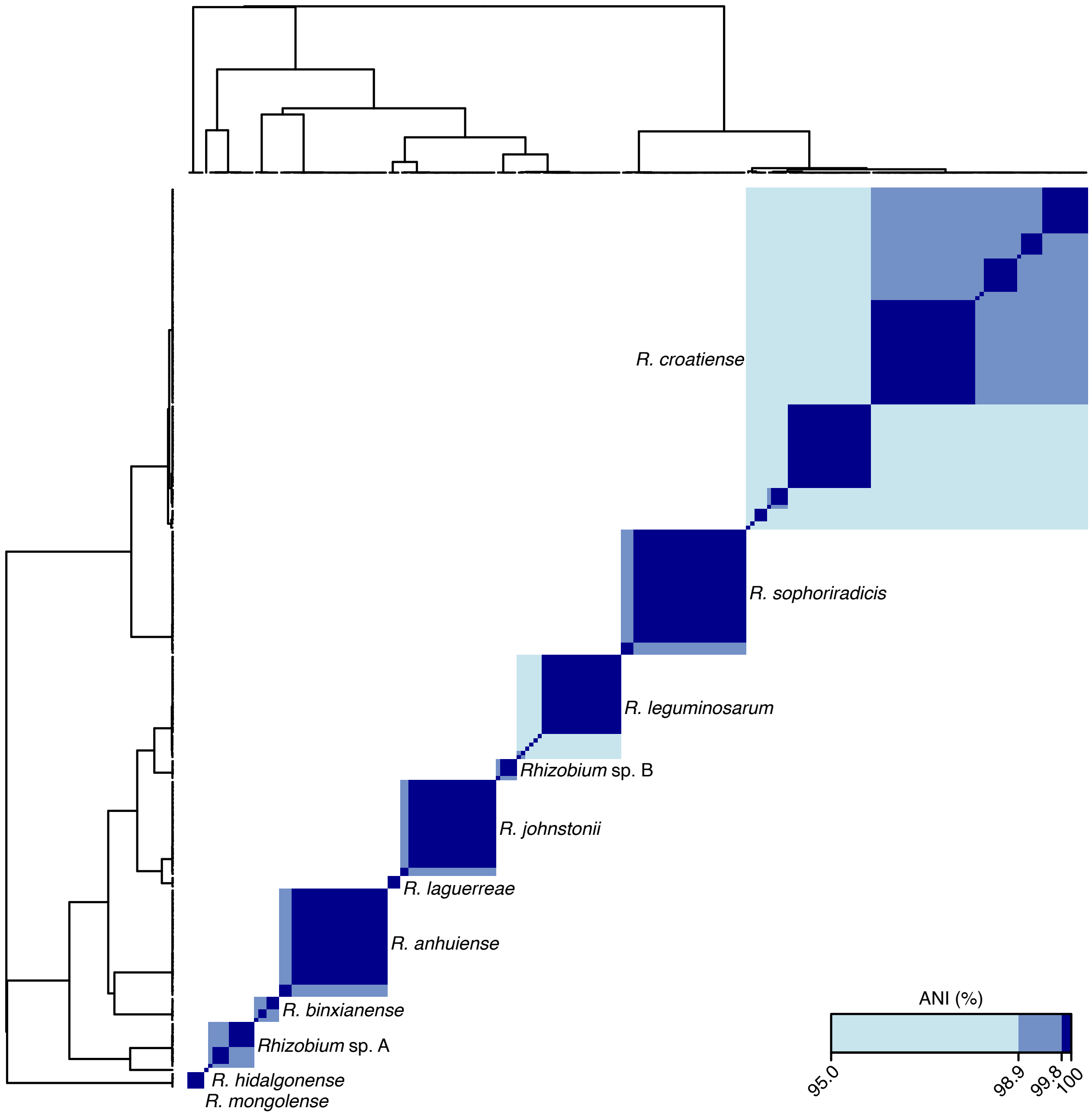


**Figure S4. Genome-based clustering of the *Rhizobium* isolates collected in this study.** The 216 *Rhizobium* isolates were clustered, and a heatma generated, based on pairwise average nucleotide identity (ANI) values calculated using the output of FastANI. Coloured squared indicated that a given pair of genomes had an ANI value ≥ 95% (light blue), ≥ 98.9% (medium blue), or ≥ 99.8% (dark blue). Individual isolates are not named as the font was too small to be legible. There are 11 clusters in which all pairwise ANI values were ≥ 95%; these represent 11 distinct species and the species name corresponding to each cluster is provided. In addition, there are 37 clusters with ANI values ≥ 99.8%, which we categorize as approximate strain-level clusters. The underlying data (with isolate names) is provided as Dataset S3.

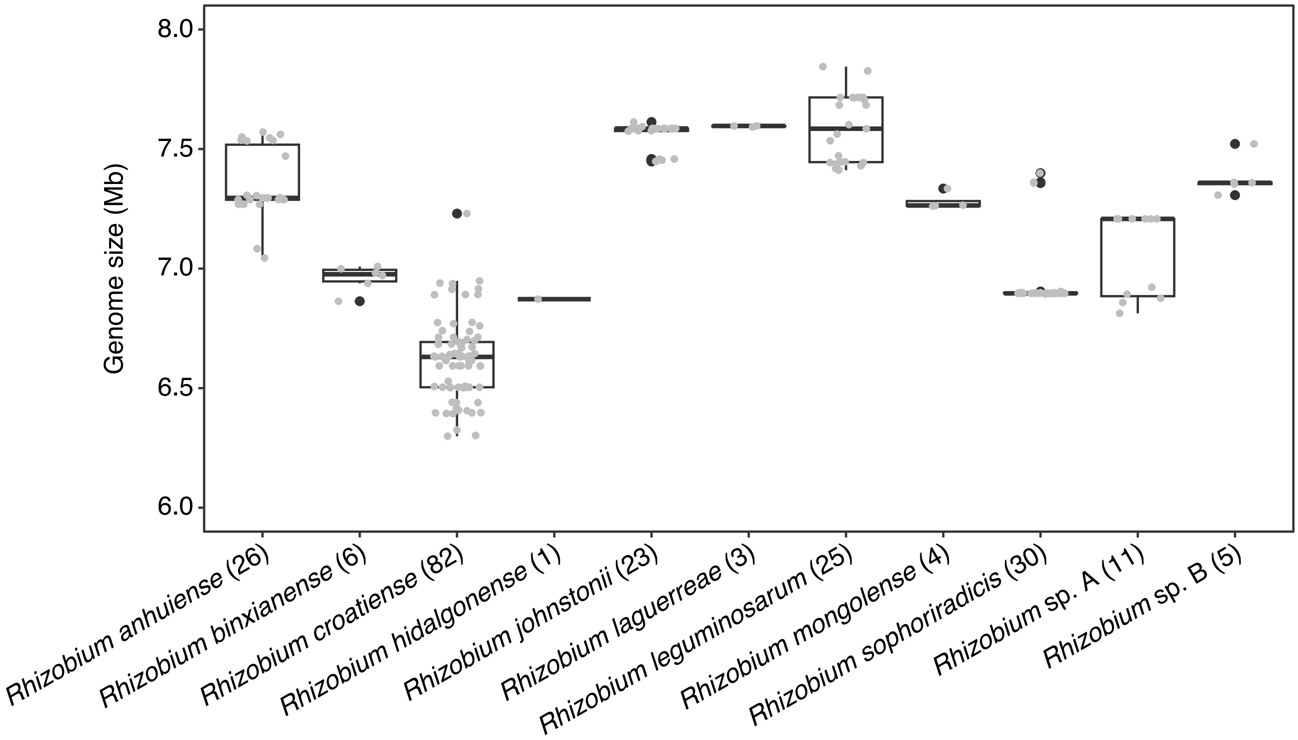


**Figure S5. Genome size distribution of *Rhizobium* isolates.** The distribution of genome sizes for each *Rhizobium* species isolated in this study is shown as box plots with individual data points overlayed as grey dots. The number of isolates in each species is given following each species name.

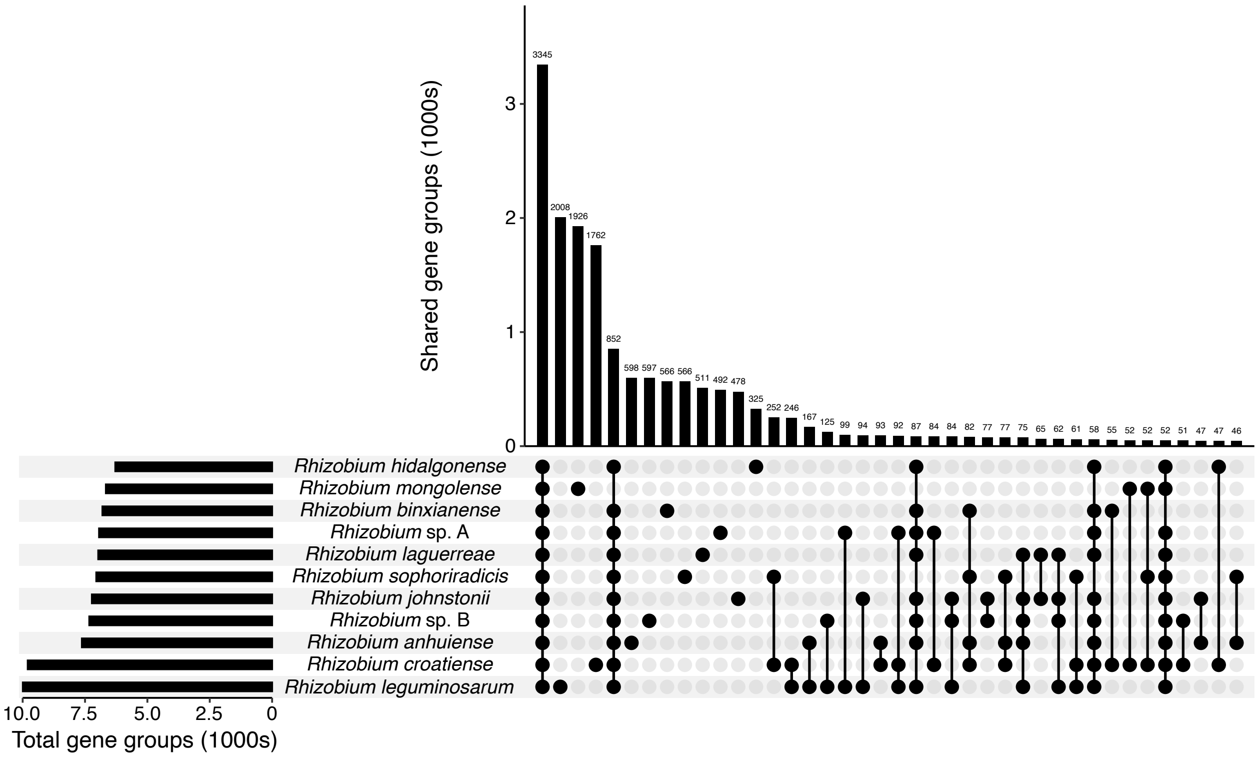


**Figure S6. Pangenome analysis of the 216 *Rhizobium* isolates.** Panaroo was used to group the genes of all 216 *Rhizobium* isolates into gene groups, after which gene presence/absence was summarized at the species level; a gene group was said to be present in a species if it was present in at least one isolate. The distribution of gene groups was then summarized using UpSetR. The bar graph on the lefthand side of the figure shows the total number of gene groups in a particular species, while the upper graph shows the number of gene groups conserved across the indicated species below the graph.

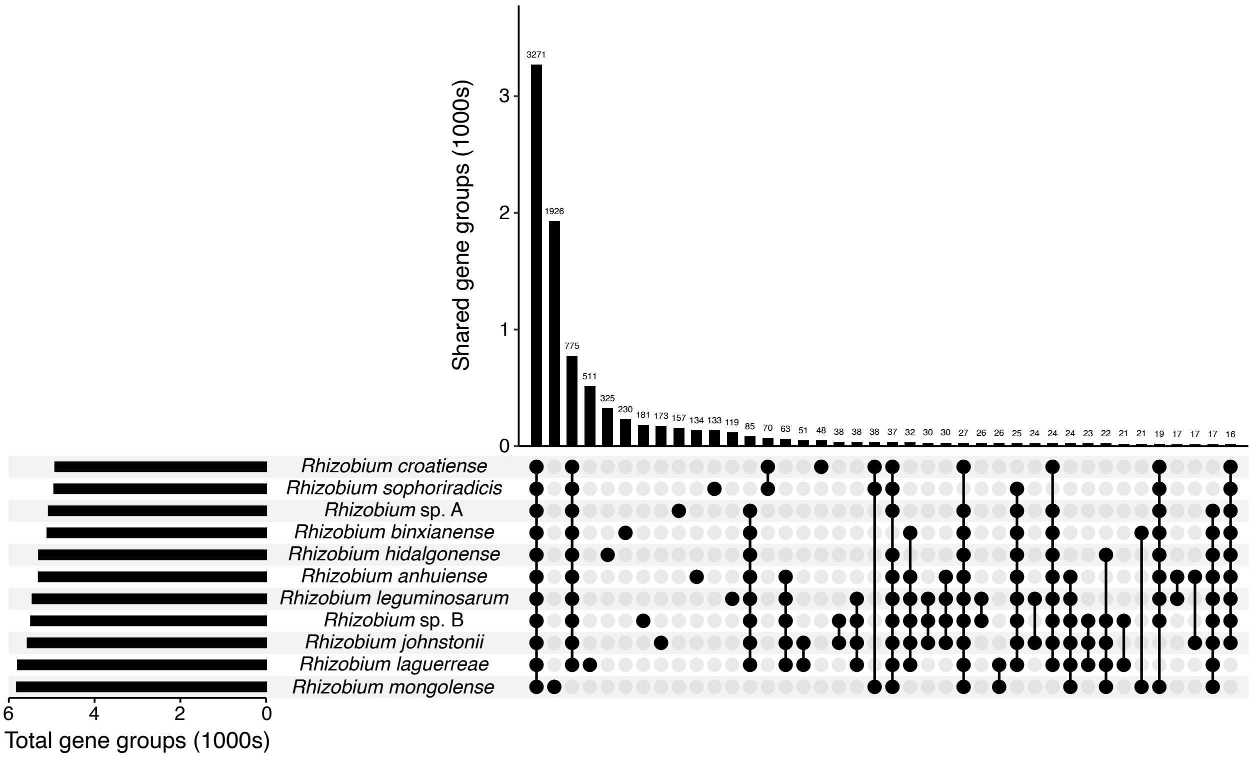


**Figure S7. Conservation of the core genome of the 11 *Rhizobium* species.** Panaroo was used to group the genes of all 216 *Rhizobium* isolates into gene groups, after which gene presence/absence was summarized at the species level. Gene groups were then filtered to only include those present in ≥ 95% of all isolates (i.e., present in the core or soft-core genome of a species) in at least one species and absent in all other species. The distribution of these gene groups was then summarized using UpSetR. The bar graph on the lefthand side of the figure shows the total number of core / soft-core gene groups in a particular species, while the upper graph shows the number of gene groups conserved across the indicated species below the graph.

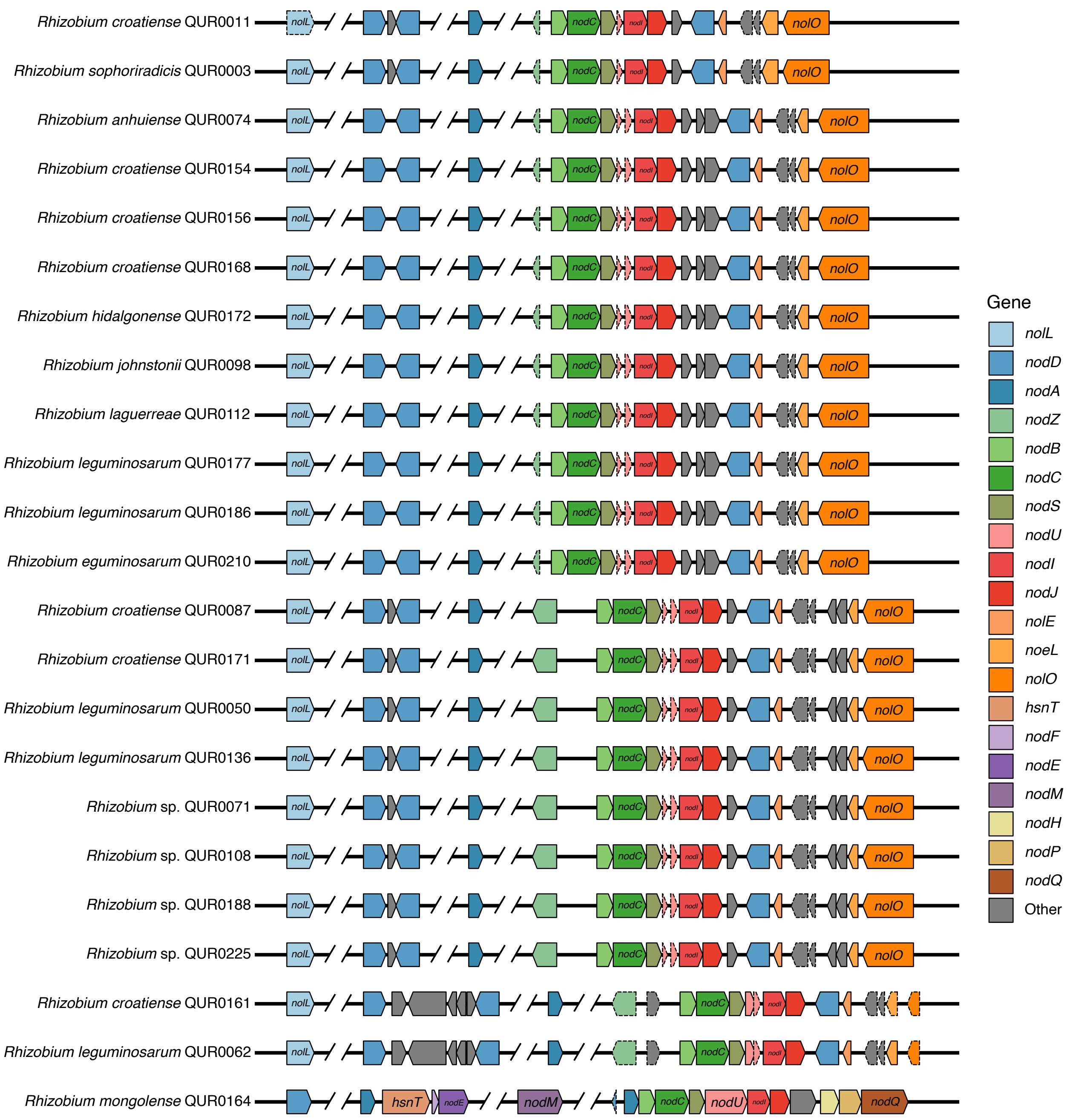


**Figure S8. Organization of *nod* genes in representative *Rhizobium* isolates.** The organization of *nod* genes was examined in 23 *Rhizobium* isolates; these included the 22 representatives of the approximate strain-level clusters that had complete genomes, as well as *Rhizobium croatiense* QUR0161 to have a second representative with the *nodC* $\gamma$-b allele. Genes are drawn to scale, orthologous genes are colour coded, and pseudogenes are indicated with dashed borders. Gaps in sequences are represented by two angled lines with white space between them.

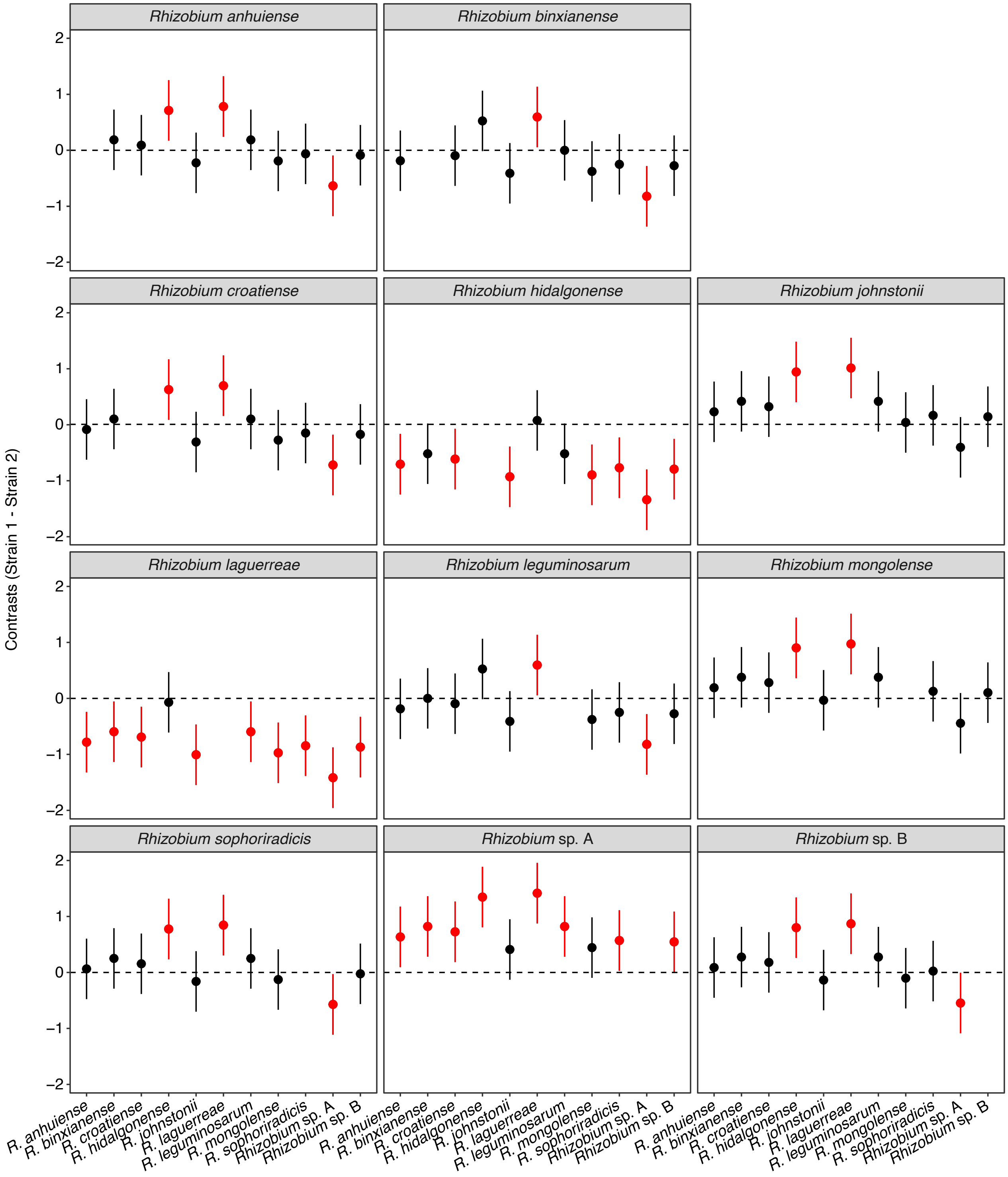


**Figure S9. Pairwise contrasts comparing the shoot dry weight of common bean plants inoculated with different *Rhizobium* species.** Pairwise contrasts were calculated using the emmeans package, by subtracting the values for plants inoculated with the species indicated along the x-axis from the values for plants inoculated with species indicated at the top of each graph. Values above 0 mean that plants inoculated with the top species were larger than those inoculated with the bottom species. Contrasts in red are statistically significant (Pr(>F) < 0.05).

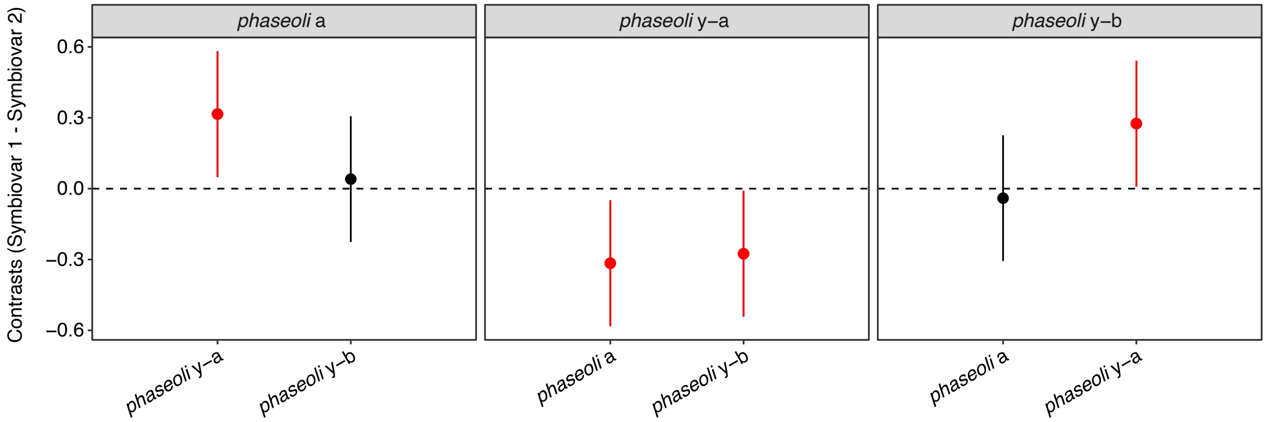


**Figure S10. Pairwise contrasts comparing the shoot dry weight of common bean plants inoculated with different *Rhizobium* symbiovar *phaseoli* subclades.** Pairwise contrasts were calculated using the emmeans package, by subtracting the values for plants inoculated with the symbiovar subclade indicated along the x-axis from the values for plants inoculated with symbiovar subclades indicated at the top of each graph. Values above 0 mean that plants inoculated with the top symbiovar subclade were larger than those inoculated with the bottom species. Contrasts in red are statistically significant (Pr(>F) < 0.05).

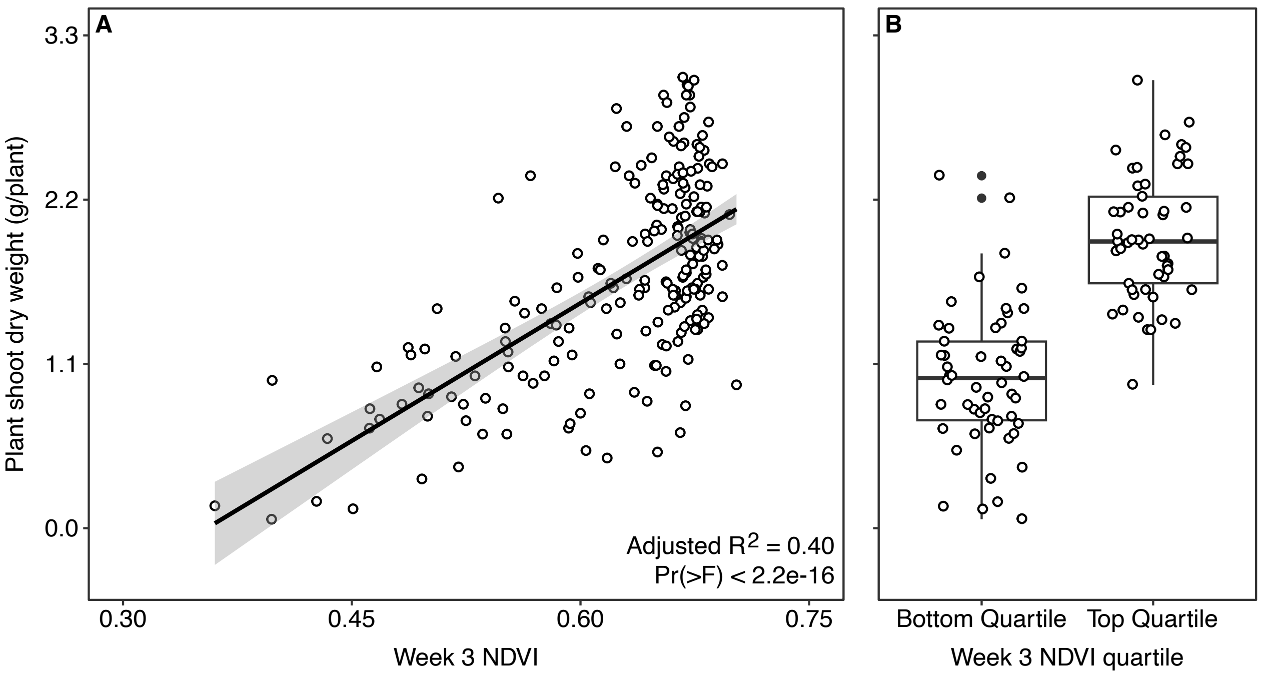


**Figure S11 – Correlation between week 3 NDVI measurements and week 5 shoot dry weights.** (**A**) A scatterplot showing the relationship between week 3 NDVI measurements and week 5 shoot dry weight measurements for navy bean plants inoculated with various *Rhizobium* isolates. Each dot represents a single plant inoculated with one of the 216 *Rhizobium* isolates, while the line represents the output of a linear model showing the relationship between the two types of measurements. (**B**) Boxplots summarizing the distribution of plant shoot dry weights for plants in the bottom quartile or top quartile of week 3 NDVI measurements. Each dot represents a single plant inoculated with a *Rhizobium* isolate, while the boxplots represent the distribution of the data.

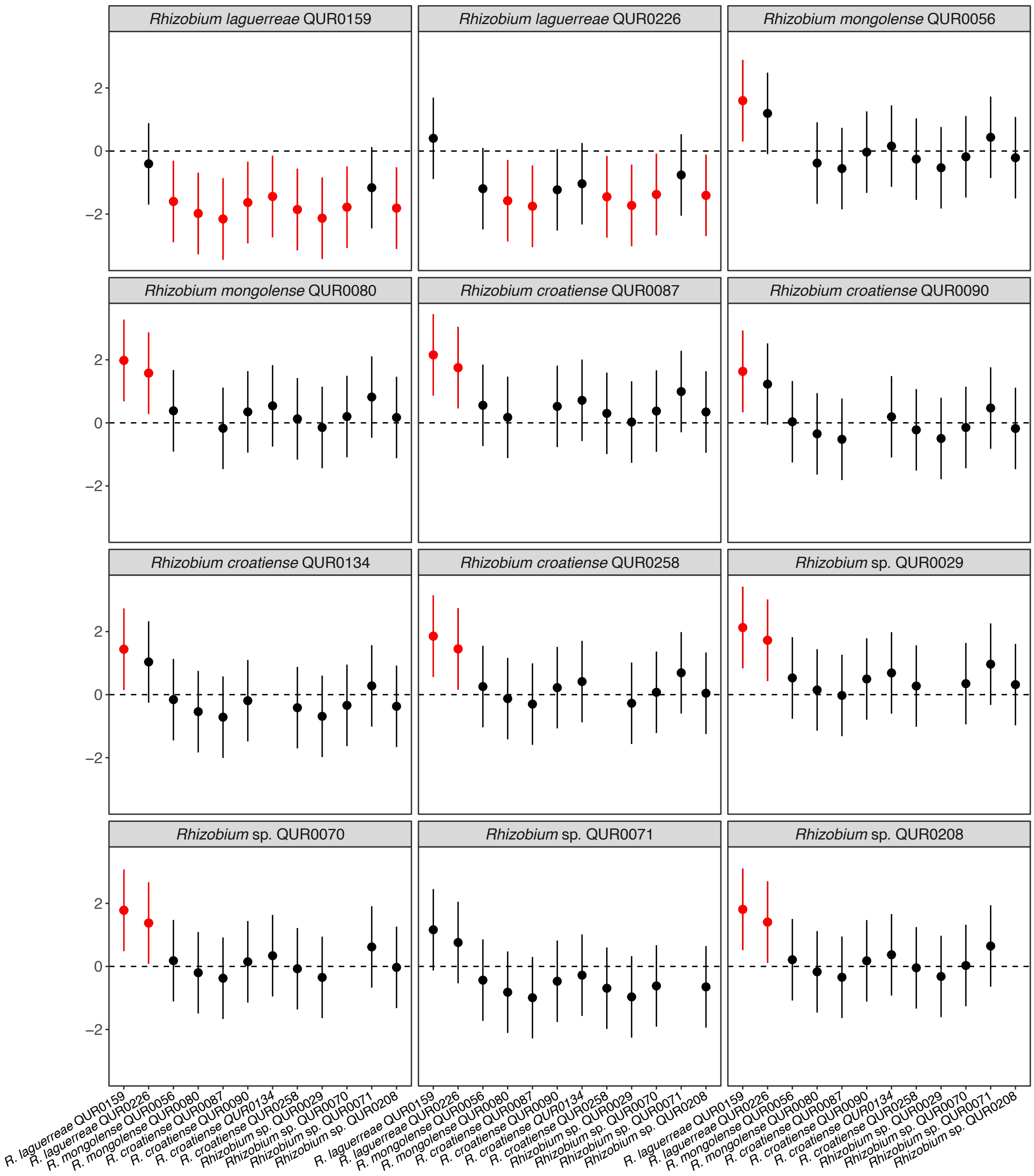


**Figure S12. Pairwise contrasts comparing the shoot dry weight of common bean plants inoculated with different *Rhizobium* isolates.** Pairwise ontrasts were calculated using the emmeans package, by subtracting the values for plants inoculated with the species indicated along the x-axis from the values for plants inoculated with species indicated at the top of each graph. Values above 0 mean that plants inoculated with the top species were larger than those inoculated with the bottom species. Contrasts in red are statistically significant (Pr(>F) < 0.05).

**Dataset S1. Isolate metadata.** Metadata is provided for each bacterial isolate collected during the nodule trapping experiment. Columns include strain name, CCASM ID, species classification, taxonomic lineage, host plant species used for nodule trapping, cultivar of host plant used for nodule trapping, soil from which the isolate was collected via nodule trapping, isolation source (nodule in all cases), year of isolation, BioProject accession for associated genomic data, assembly status (complete or draft), genome size, number of contigs in the assembly, number of protein coding genes, number of pseudogenes, and genome completeness and contamination as determined with CheckM.

**Dataset S2. *Rhizobium* and *Martinezella* type strains.** The species name and strain name of the *Rhizobium* and *Martinezella* type strains used in the phylogenetic analysis. In addition, accession numbers are provided to download the corresponding genomes from NCBI (accession starts with GCF_), ENA (accession starts with ERZ), or JGI GOLD (accession starts with Ga).

**Dataset S3. Pairwise average nucleotide identity (ANI) values.** A matrix containing the pairwise ANI values calculated between each pair of genomes for all 216 *Rhizobium* isolates collected in this study.

**Dataset S4. Plant shoot dry weights ad NDVI measurements from theinitial screen of *Rhizobium* isolates.** The shoot dry weight and weekly NDVI measures are provided for plants inoculated with 216 *Rhizobium* isolates as well as controls without rhizobia and either with (Nitrogen_plus) or without (Nitrogen_negative) nitrogen supplementation.

**Dataset S5. Plant shoot dry weights from follow-up screen of a subset of *Rhizobium* isolates.** Shoot dry weights are provided for all replicate plants inoculated with 12 *Rhizobium* isolates, as well as uninoculated controls.

**Dataset S6. Plant shoot dry weights from the nitrogen addition experiment.** Shoot dry weights are provided for all replicate plants with or without *Rhizobium* inoculation under different nitrogen regimens.
